# Species-specific differences in antagonism of APOBEC3 proteins by HIV-2 and SIVsmm Vif proteins

**DOI:** 10.1101/2020.09.08.287177

**Authors:** Rayhane Nchioua, Dorota Kmiec, Amit Gaba, Christina M. Stürzel, Tyson Follack, Stephen Patrick, Andrea Kirmaier, Welkin E. Johnson, Beatrice H. Hahn, Linda Chelico, Frank Kirchhoff

## Abstract

SIVsmm infecting sooty mangabeys has been transmitted to humans on at least nine independent occasions, giving rise to HIV-2 groups A to I. SIVsmm isolates replicate in human T cells and seem capable of overcoming major human restriction factors without adaptation. However, only groups A and B are responsible for the HIV-2 epidemic in Sub-Saharan Africa and it is largely unclear whether adaptive changes were associated with significant spread in humans. To address this, we examined the sensitivity of infectious molecular clones (IMCs) of five HIV-2 strains (4 group A and one AB recombinant) and representatives of five different SIVsmm lineages to inhibition by type I interferon (IFN) and various APOBEC3 proteins. We confirmed that SIVsmm strains replicate in primary human CD4+ T cells. However, SIVsmm replication was highly variable, typically lower relative to HIV-2 isolates and almost entirely prevented by type I IFN treatment. Viral propagation was generally dependent on intact *vif* genes, highlighting the need for efficient counteraction of APOBEC3 proteins. On average, SIVsmm strains were significantly more susceptible to inhibition by human APOBEC3D, F, G and H than HIV-2 IMCs. For example, human APOBEC3F reduced infectious virus yield of SIVsmm by ∼80% but achieved only ∼40% in the case of HIV-2. Functional and mutational analyses of human, sooty mangabey and rhesus macaque derived alleles revealed that an R128T polymorphism in APOBEC3F is important for species-specific counteraction by HIV-2 and SIVsmm Vif proteins. In addition, we found that changes of Y45H and T84S in SIVsmm Vif increase its ability to antagonize human APOBEC3F. Altogether, our results show that SIVsmm Vifs show some intrinsic activity against human ABOBEC3 proteins, but HIV-2 Vifs acquired adaptive changes to efficiently clear this barrier in the human host.

**AUTHOR SUMMARY:** SIVs infecting African monkey species do not infect humans, with one notable exception. SIVsmm from sooty mangabeys managed to cross the species barrier to humans on at least nine independent occasions. This is because SIVsmm strains seem capable of overcoming many innate defense mechanisms without adaptation and that their Vif proteins are active against human APOBEC3 proteins. Here, we show that replication of SIVsmm is highly variable in human CD4 T cells and more sensitive to interferon inhibition compared to HIV-2. While different lineages of SIVsmm were capable of counteracting human APOBEC3 proteins in a Vif-dependent manner, they were significantly more susceptible to inhibition by APOBEC3D/F/G/H compared to HIV-2. Mutational analyses revealed an R128T substitution in APOBEC3F and a T84S change in Vif are relevant for species-specific counteraction by HIV-2 and SIVsmm. Altogether, our results support that HIV-2 group A adapted to humans prior to or during epidemic spread.

## INTRODUCTION

Simian immunodeficiency viruses (SIV) have been infecting primates for many hundreds of thousands or even millions of years [1]. However, only in the 20th century, cross-species transmissions from three of more than forty non-human primate species harbouring SIVs gave rise to Human Immunodeficiency Virus (HIV) [1,2]. There are two types of the virus, HIV-1 and HIV-2, which are further subdivided into four and nine groups, respectively, each originating from an independent transmission from great apes or monkeys. HIV-1 group M (major), which is responsible for at least 95% of all infections, as well as HIV-1 group N that has only been detected in about 20 individuals, originated from SIVcpz infecting chimpanzees [1,2]. SIVcpz was also transmitted from chimpanzees to gorillas, which passed SIVgor to humans on two occasions and gave rise to epidemic HIV-1 group O strains and the very rare group P of HIV-1 [2].

It is plausible that great apes and not monkeys have transmitted their viruses to humans since these *Homininae* are genetically closely related. However, there is a striking exception. SIVsmm infecting sooty mangabeys has been transmitted to humans on at least nine independent occasions giving rise to HIV-2 groups A-I [2,3]. However, only two of the nine groups of HIV-2 (A and B) have spread significantly in the human population and are responsible for about one to two million infections, mostly in West Africa [4]. SIVsmm has also been accidentally transmitted to rhesus macaques in primate centres and SIVmac infection of Asian macaques became a valuable animal model for HIV pathogenesis and AIDS in humans [5].

Several factors help to explain why SIVsmm from sooty mangabeys frequently crossed the species barrier to humans, while no other SIVs infecting numerous monkey species in sub-Saharan Africa have been detected in humans. SIVsmm is highly prevalent in sooty mangabeys and these monkeys are kept as household pets or hunted for bushmeat suggesting frequent viral exposure of humans. In addition, SIVsmm isolates replicate in human PBMCs [6] and seem capable of counteracting some major human restriction factors without adaptation [7]. For example, the SIVsmm Env is active against human tetherin [8,9] and the SIVsmm Vpx protein counteracts the human ortholog of SAMHD1 [10] as well as the human HUSH complex [11]. In addition, human SERINC5 does not pose a barrier to zoonotic transmission as it is counteracted by SIVsmm Nef [12]. Altogether, SIVsmm accessory proteins seem to be more active against human restriction factors than those of SIVs infecting other monkey species.

Apolipoprotein B mRNA editing enzyme catalytic polypeptide-like 3 (APOBEC3; A3) represent perhaps the best studied and most relevant antiretroviral restriction factors [13,14]. These cellular cytidine deaminases bind viral RNA, are encapsidated into newly formed virions and impair viral infectivity by inducing the formation of deleterious G-to-A hyper-mutations in the viral genome and by interfering with reverse transcription [15–17]. Humans possess seven A3 proteins (A, B, C, D, F, G and H) that resulted from several gene duplications on chromosome 22 [18]. It has been reported that all human A3 proteins, except A3A, have the capacity to suppress HIV-1 group M infectivity in CD4+ T cells [19]. A3A may restrict HIV infectivity in monocyte derived cells [20,21]. However, A3B can only suppress HIV infectivity when expressed and encapsidated into HIV virions from 293T producer cells, which is not relevant *in vivo* where it is localized in the nucleus and unable to encapsidate into virions [22]. A3C is highly expressed in CD4+ T cells, but hardly able to induce enough mutations in the HIV proviral DNA to affect infectivity [23]. Primate lentiviruses use their Vif protein to antagonize restriction by A3 proteins. All lentiviral Vif proteins interact with A3 proteins to recruit them to the Cullin 5/ElonginBC E3 ubiquitin ligase complex for proteasomal degradation [15]. The importance of this viral counteraction mechanism is evident from the fact that Vif is also found in feline immunodeficiency viruses (FIVs) and already present in prosimian lentiviruses that entered the germline of lemurs about 4 million year ago [17,24]. In addition, a functional *vif* gene is critical for replication of SIVmac in rhesus macaques [25]. The ability of primate lentiviral Vif proteins to promote infectivity is species-specific [26]. Notably, it has been shown that the recombination events between SIVs originating from small monkeys that lead to the emergence of SIVcpz in chimpanzees resulted in a functional Vif protein at the cost of the accessory *vpx* gene [13]. The adaptive changes allowing the precursors of HIV-1 to counteract A3 proteins and other restricting factors after crossing the species barriers first from monkeys to chimpanzees and later from great apes to humans have been well studied [13,27–30].

It has been reported that SIVsmm Vif proteins show some activity against APOBEC3G from many species including the human homolog [29]. However, counteraction of human A3 proteins may be imperfect since epidemic HIV-2 strains show on average more APOBEC3G/F-induced hyper-mutations than HIV-1 isolates [31]. It is incompletely understood how susceptible SIVsmm is to IFN-inducible factors in human cells and whether their Vif proteins acquired adaptive changes allowing more effective counteraction of A3 proteins following zoonotic transmission to humans. Here, we show that SIVsmm has lower replication fitness than HIV-2 in primary human cells, especially in the presence of IFN. We further demonstrate that SIVsmm Vifs show some activity against human ABOBEC3 proteins but HIV-2 Vifs acquired adaptive changes to efficiently overcome this barrier in the human host.

## RESULTS

### Replication fitness and IFNα sensitivity of HIV-1, HIV-2 and SIVsmm in CD4+ T cells

Primary SIVsmm isolates are capable of replicating in human cells without adaptive changes [6]. However, the replication kinetics of SIVsmm and HIV-2 in human CD4+ T cells and their susceptibility to the inhibitory effects of type I IFNs have not been directly compared. Thus, it remained largely unclear whether HIV-2 acquired increased replication fitness in human cells and reduced susceptibility to IFN-inducible antiviral factors during adaptation to the human host. To address this, we analyzed a panel of five infectious molecular clones (IMCs) representing epidemic (A and CRF01_AB) groups of HIV-2, as well as six SIVsmm IMCs representing five different lineages (Table S1). For comparison, we examined four transmitted-founder (TF) HIV-1 strains [32,33] and the T-cell line adapted HIV-1 NL4-3 IMC, as well as SIVmac239 [34], a macaque-passaged derivative of SIVsmm, that has been commonly used in non-human primate studies on viral pathogenesis and vaccine development.

To determine the replicative capacity of HIV-1, HIV-2 and SIVsmm, we infected activated human peripheral blood mononuclear cells (PBMCs) with virus stocks normalized for reverse transcriptase (RT) activity and determined the infectious virus yield in culture supernatants at various days post-infection by TZM-bl reporter cell infection assay. All ten HIV-1 and HIV-2 IMCs replicated efficiently in human PBMCs (Figure 1A). On average, the HIV-2 IMCs achieved lower infectious virus yields than HIV-1 IMCs (Figure 1B). However, the HIV-2 GH123 construct derived from an AIDS patient from Ghana [35] displayed surprisingly rapid kinetics and achieved ∼10-fold higher infectious virus yields than the remaining four HIV-2 IMCs (Figure 1A). Notably, this HIV-2 strain has a truncated gp41 cytoplasmic tail. The RT activities in the supernatant of PBMCs infected with HIV-2 GH123 were similar to those obtained for other HIV-2 IMCs (data not shown) indicating that the gp41 truncation increases viral infectivity for TZM-bl reporter cells as previously reported for SIVmac239 [36] and HIV-2 ST in SupT1 cells [37].

**Figure 1:**
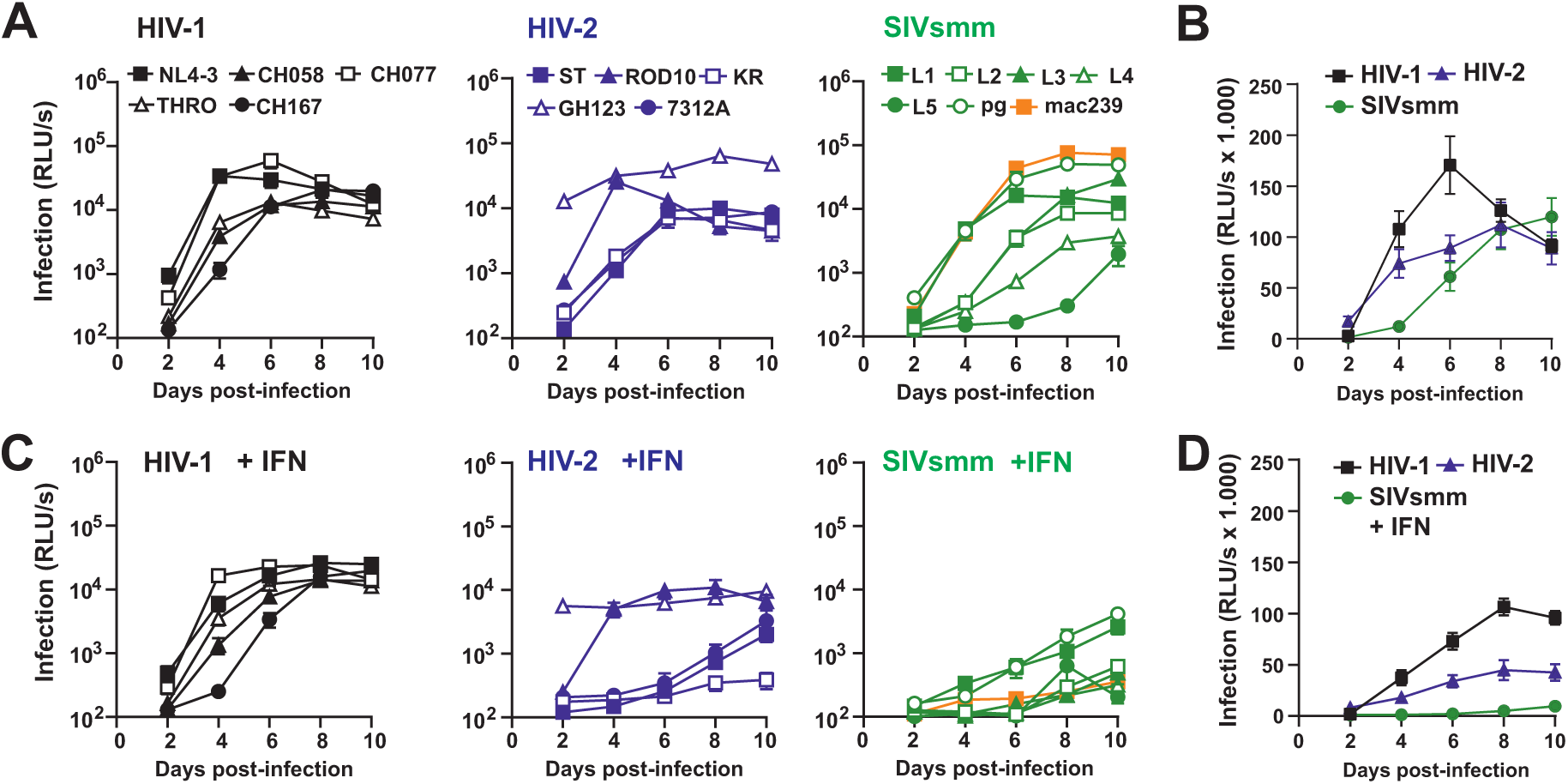
HIV-1, HIV-2 and SIVsmm differ significantly in IFN sensitivity. (**A**) Replication of HIV-1, HIV-2 and SIV proviral constructs in primary human CD4+ T cells. The results show mean infectious virus yields (n = 3) measured by infection of TZM-bl reporter cells with normalized volumes of the supernatants infected CD4+ T cell cultures derived from three different PBMC donors. (**B**) Mean infectious virus yield (±SEM) of the five HIV-1, five HIV-2 and six SIVsmm IMCs measured at the indicated days post-infection. (**C**) Replication of HIV-1, HIV-2 and SIV IMCs in CD4+ T cells infected as described in panel A the presence of IFN-α. (**D**) Mean infectious virus yields of HIV-1, HIV-2 and SIVsmm detected in the presence of IFN-α.

The replicative capacity of the SIV constructs in primary human cells was more variable compared to HIV-1 and HIV-2. The SIVmac239 and SIVsmm PG strains replicated as efficiently as HIV-1 strains (Figure 1A). Notably, SIVmac239 was passaged in human HUT-78 cells [34], and SIVsmm PG was passaged in human PBMCs and CEMx174 cells after isolation from lymph nodes of an infected pig-tailed macaque [38]. Thus, their effective replication may reflect adaptation for growth in human cells. The remaining SIVsmm clones represent five divergent lineages [39] and were obtained after passage in rhesus macaques that are genetically closely related to sooty mangabeys (Table S1) [6]. Like HIV-2, the SIVsmm lineage 1 (L1) strain replicated efficiently in human PBMCs, while the SIVsmm L5 construct showed only marginal levels of replication (Figure 1A). The remaining three IMCs (L2, L3 and L4) displayed a phenotype intermediate between SIVsmm L1 and L5. On average, the HIV-1 IMCs replicated with the highest efficiency in human CD4+ T cells and HIV-2 clones showed more rapid replication kinetics than SIVsmm IMCs (Figure 1B).

Type I IFN induces numerous antiviral factors that are frequently counteracted by primate lentiviruses in a species-specific manner [40–42]. Previous studies showed that SIVsmm strains are capable of antagonizing several major human restriction factors [7]. It is poorly understood, however, whether they counteract the antiviral effects of IFN in primary human T cells as effectively as HIV-2. To address this, we infected activated human PBMCs in the presence of IFN-α. We found that treatment with IFN-α (500 U/ml) reduced infectious yield of HIV-1 about 2-fold and resulted in moderately delayed replication kinetics (Figure 1C). In comparison, only the HIV-2 GH123 and ROD10 strains replicated efficiently in the presence of IFN-α, while infectious virus production was grossly impaired for 7312A and ST and essentially absent for KR (Figure 1C). The inhibitory effect of IFN-α on SIVsmm replication was even more severe and only the SIVsmm PG and L1 clones replicated to clearly detectable levels (Figure 1C). The SIVmac239 IMC that showed the highest infectious virus yield in the absence of IFN-α was almost completely suppressed in its presence. Altogether, the results showed that HIV-1 IMCs display higher replication fitness in primary human CD4+ T cells than HIV-2 IMCs in both the absence (Figure 1B) and presence of IFN-α (Figure 1D). On average, however, HIV-2 was significantly more competent for replication in primary human cells than SIVsmm, indicating significant adaptation for effective spread as well as counteraction of IFN-inducible innate defense factors in the human host.

### HIV-2 is less susceptible to inhibition by human APOBEC3 proteins than SIVsmm

Our results indicated that during the emergence of HIV-2 from SIVsmm, the virus adapted to become more fit for replication in the human host. To define underlying mechanisms, we investigated the potential role of Vif-mediated antagonism of APOBEC3 proteins in the evolution of replication fitness in human cells in the SIVsmm/HIV-2 lineage. To verify the importance of Vif for effective viral replication, we generated *vif*-defective derivatives of six HIV and SIV IMCs. The *vif* defective HIV-1 NL4-3, CH077, CH058 as well as HIV-2 7312A constructs showed strongly delayed and severely attenuated replication kinetics in both the absence and presence of IFN-α (Figure 2A). A defective *vif* gene further impaired the already modest replication of SIVsmm L5. Unexpectedly, the *vif*-defective SIVmac239 construct showed higher levels of replication than *vif*-defective HIV-1 and HIV-2 IMCs in the absence of IFN-α (Figure 2A). Predictably, these analyses verified that functional Vif expression is required for effective lentiviral replication in primary human T cells.

**Figure 2:**
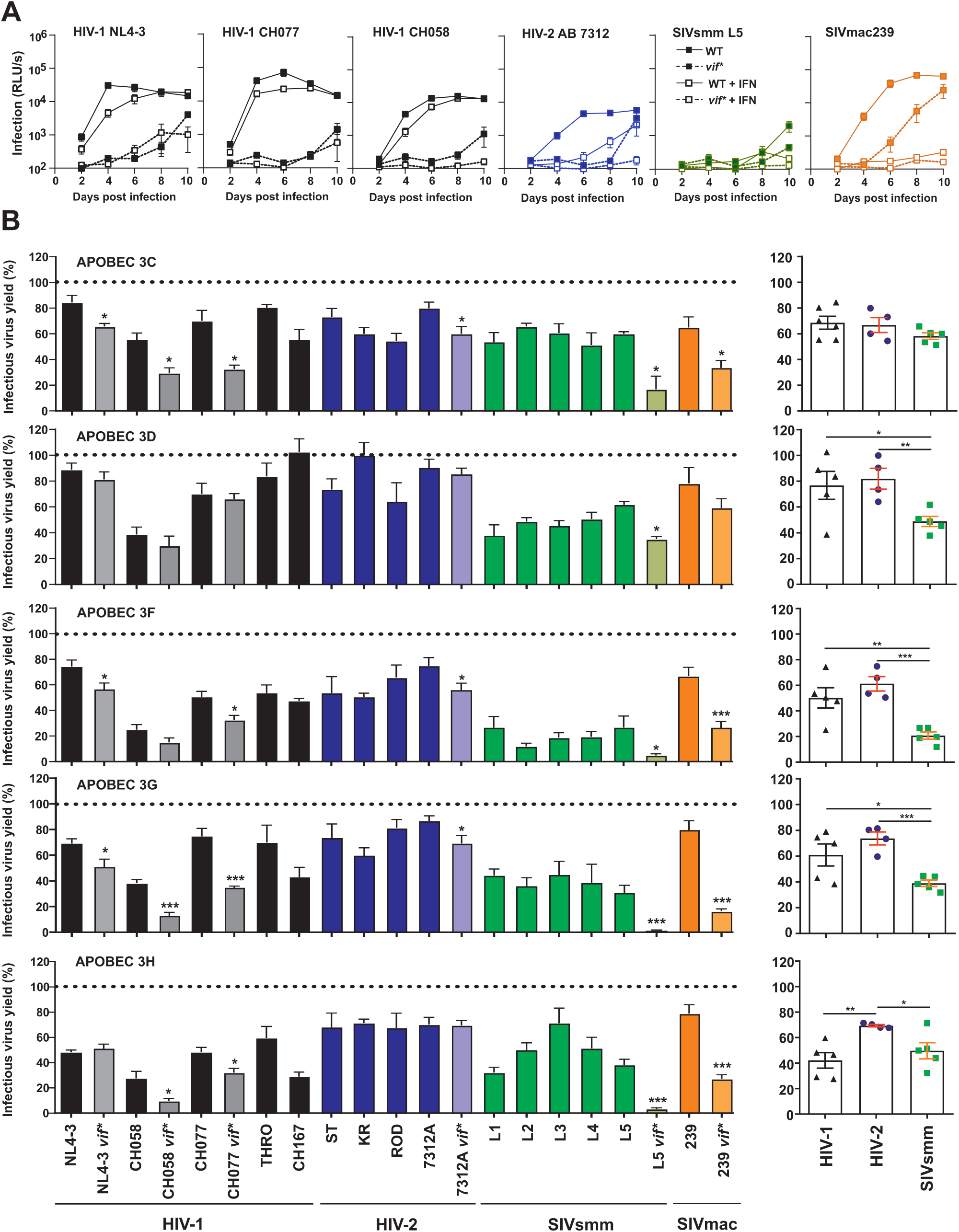
Sensitivity of HIV-1, HIV-2 and SIVsmm to human APOBEC3 family members. (A) Relevance of an intact *vif* gene for HIV-1, HIV-2 and SIV replication in primary human CD4+ T cells. Replication kinetics of HIV-1, HIV-2 or SIVsmm IMCs containing intact or disrupted vif genes in CD4+ T cells in the presence of 500 U/ml IFN-α (dashed lines) or absence of IFN-α (solid lines). Infectious virus yield was measured using the TZM-bl reporter cell infectivity assay. (**B**) Proviral constructs of the indicated IMCs of HIV-1, HIV-2, or SIVsmm and a plasmid expressing the various APOBEC3 proteins were cotransfected into HEK293T cells. Infectious virus yield was measured using the TZM-bl reporter cell infectivity assay. For each proviral construct, values were normalized to the infectious virus yield obtained in the absence of APOBEC3 expression construct (100%). Shown is the mean from 3 to 8 independent experiments each measured in triplicates ± SEM. The right panel shows comparisons of the sensitivity of the HIV-1, HIV-2 and SIVsmm group to the various APOBEC3 family members. Symbols represent the average infectious virus yield relative to the absence of APOBEC3s (100%) obtained for individual IMCs. Bar diagram shows the infectious virus yield obtained for all IMCs from the indicated groups. *, P < 0.05; **, P < 0.01; ***, P < 0.001, calculated using Student’s t test.

To determine possible differences in the susceptibility of HIV-1, HIV-2 and SIVsmm to various human APOBEC3 proteins, we measured infectious virus yield from HEK293T cells following cotransfection of the proviral constructs with expression vectors for various APOBEC3s or an empty control vector (Figures 2B, S1). This system was chosen due to the reported lack of or only low expression of endogenous APOBEC3 genes in this cell line. In agreement with these reports, we observed no infectivity defects of *vif\** mutant IMCs in the absence of APOBEC3 overexpression (Figure S1). All five APOBEC3 proteins analyzed (C, D, F, G and H haplotype II) inhibited infectious HIV-1 production to some extent with average efficiencies ranging from 20% (D) to 60% (H) (Figure 2B). As expected, *vif*-defective HIV and SIV IMCs were usually more susceptible to APOBEC3 inhibition than the parental WT viruses. However, NL4-3 Vif was less effective in counteracting A3G than the Vif proteins of the primary HIV-1 CH058 strain. In addition, the effects of intact *vif* genes on infectious virus yield were usually modest in the case of A3C, A3D and A3F, most likely due to both relatively low antiviral activity and ineffective counteraction by Vif. Surprisingly, the *vif*-defective HIV-2 7312A construct was largely resistant to all APOBEC3 proteins. In contrast, lack of Vif function generally increased the susceptibility of SIVsmm L5 and SIVmac239 especially to human A3F, A3G and A3H (Figure 2B). Altogether, the susceptibility of HIV-2 strains did not differ significantly from those of HIV-1 for APOBEC 3C, 3D, 3F and 3G (Figure 2B). Unexpectedly, HIV-2 IMCs were less sensitive to inhibition by A3H than HIV-1 (Figure 2B, bottom). Most notably, SIVsmm IMCs were on average significantly more susceptible to inhibition by human A3D, A3F, A3G and A3H than HIV-2 (Figure 2B). In contrast, SIVmac239 was largely resistant to all human APOBEC3 proteins investigated. Altogether, these results suggest that HIV-2 strains acquired changes increasing their ability to counteract human APOBEC3D, F, G and H proteins after zoonotic transmission of SIVsmm from sooty mangabeys to humans. Unexpectedly, *vif*-defective HIV and SIV IMCs differ substantially in their susceptibility to human APOBEC3 proteins. Notably, these differences were not just due to differences in the absolute levels of infectious virus production (Figure S1). Thus, although Vif function clearly plays a key role in primate lentiviral susceptibility to APOBEC3 proteins, alternative mechanisms also seem to be involved.

### HIV-2 counteracts human APOBEC3 proteins more efficiently than SIVsmm

To determine whether the reduced susceptibility of HIV-2 to human A3D, A3F, A3G and A3H relative to SIVsmm is associated with differences in the ability of the respective Vif proteins to induce APOBEC3 degradation, we performed immunoblot analyses. We cotransfected HEK293T cells with proviral HIV-1, HIV-2 or SIV and APOBEC expression constructs and measured the levels of viral and APOBEC3 proteins in the cellular extracts and culture supernatants as well as the corresponding infectious virus yields. HIV-1, HIV-2 and SIVsmm constructs are highly divergent and there are relatively few reagents specifically designed to study the latter two. Thus, antibody recognition of the p24 and p27 capsid antigens varied. Therefore, we focused on the levels of cell-associated APOBEC3 antigens normalized for cellular GAPDH or tubulin levels for quantitative comparisons. In agreement with the results on infectious virus yields (Figure 2B), the levels of cell-associated A3D, A3F, A3G and A3H were significantly lower in cells transfected with HIV-2 constructs compared to those that received SIVsmm (Figures 3A, 3B, S2A). As expected, lack of intact *vif* genes increased APOBEC3 steady-state expression levels. It came as surprise, however, that the ability of HIV-1 IMCs to degrade A3D, A3F, A3G and A3H proteins varied substantially and was usually less effective compared to HIV-2 IMCs (Figure S2A). Altogether, the levels of A3F and A3G expression relative to GAPDH correlated with the infectious virus yields in the cell culture supernatant albeit with substantial variation (Figure 3A, 3B, right panels). In agreement with previous data [43], the levels of APOBEC3 proteins in cellular extract did not always match those detectable in the viral supernatants (Figures 3A, 3B, S2A). Thus, while Vif-mediated degradation and cellular expression levels of APOBEC3 proteins are a major determinant of infectious virus production, not all HIV-1, HIV-2 and SIV constructs seem to be equally susceptible to the inhibitory effect of these APOBEC3 proteins. In addition, our results add to the evidence that Vif might also counteract APOBEC3 proteins by degradation-independent activities [44].

**Figure 3:**
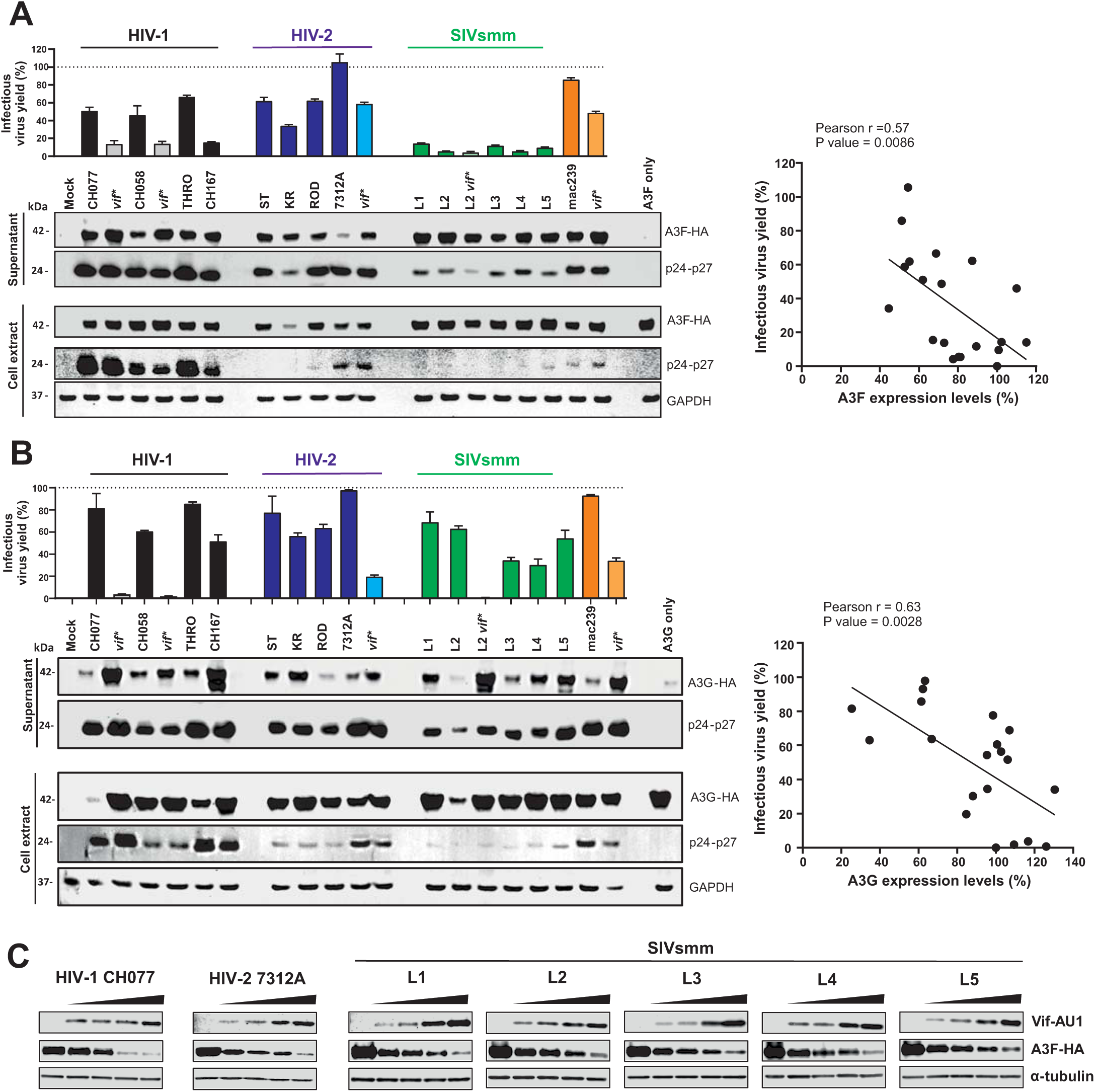
Effect of HIV-1, HIV-2 and SIVsmm on A3F and A3G protein expression levels. (**A, B**) Infectious HIV-1 yield (top) and expression levels of viral proteins and human (A) A3F or (B) A3G in HEK293T cells transfected with the indicated proviral constructs. The right panels show the correlations between infectious virus yield and the A3F or A3G protein expression levels in cellular extracts measured in the presence of proviral constructs relative to those measured in the presence of the control vector. All A3F and A3G expression values were normalized for the GAPDH loading control. Infectious virus yield was measured using the TZM-bl reporter cell infectivity assay and values were normalized to those obtained in the absence of APOBEC3 expression construct (100%). Shown are mean values (±SD) from triplicate infection. (**C**) Vif mediated degradation of A3F. 293T cells were cotransfected with constant quantity of A3F expression plasmid (100ng) and increasing amounts of HIV or SIV Vif-AU1 expression constructs. Forty-eight hours post transfection, cell lysates were collected. Proteins from cell lysates were separated by SDS PAGE, transferred to nitrocellulose membrane and probed using antibodies, as labeled, with α-tubulin serving as the loading control.

Human A3F was detected at relatively high levels in SIVsmm particles and not efficiently counteracted by SIVsmm Vif (Figure 3A). To investigate this further, we cotransfected 293T cells with constant quantities of A3F or A3G expression plasmids and increasing amounts of HIV or SIV Vif constructs. All Vif proteins analyzed induced efficient degradation of A3G, while none of them efficiently counteracted A3F at high expression levels (Figure S2B). To detect more subtle activities of HIV and SIV Vif proteins in degrading human A3F, we performed the experiment using a 5-fold lower dose of A3F expression constructs. Under these conditions, a dose-dependent reduction of A3F expression was detectable for all Vif proteins examined (Figure 3C). In agreement with the results obtained using the proviral constructs (Figure 3A), the HIV-1 CH077 and HIV-2 7312A Vifs degraded A3F more efficiently than the SIVsmm Vifs (Figure 3C). Altogether, these results show that HIV-2 acquired an increased ability to degrade various human APOBEC3 proteins and further suggest that especially A3F might represent a barrier to successful spread of SIVsmm after zoonotic transmission.

### HIV-2 and SIVsmm show species-specific differences in A3 antagonism

To further examine the species-specific evolution of APOBEC3 antagonism in the HIV-2/SIVsmm/mac lineage, we compared the susceptibility of HIV-2 and SIVsmm to inhibition by human and monkey-derived A3F, A3G and A3H orthologues. Two A3F alleles, each, were available from sooty mangabeys and macaques. The two SMM A3F alleles differed in four amino acid residues from one another, in 14 residues from the macaque homologue and in ∼50 residues from the human version (Figure 4A). Most variations did not fall within previously proposed active site residues, and the two Zinc-coordinating Cys residues as well as the catalytic Glu residues are generally conserved. Functional analyses showed that HIV-1 CH077 efficiently antagonizes human and sooty mangabey A3Fs in a Vif-dependent manner but is inhibited by ∼50% by the macaque orthologues (Figure 4B, 4C). In comparison, the HIV-2 ST, ROD and 7312A IMCs as well as SIVmac239 were resistant to A3F orthologues from all three species investigated (Figure 4B). With the single exception of SIVsmm L5 that shows unique variations at amino acid positions 93, 134, 156 and 168 of Vif, all four SIVsmm IMCs analyzed were highly susceptible to human A3F but resistant to inhibition by the sooty mangabey and macaque A3Fs (Figure 4B). On average, HIV-2 was significantly less sensitive to human A3F than SIVsmm (Figure 4C) suggesting human-specific adaptation or zoonotic transmission of a pre-adapted, less susceptible SIVsmm variant.

**Figure 4:**
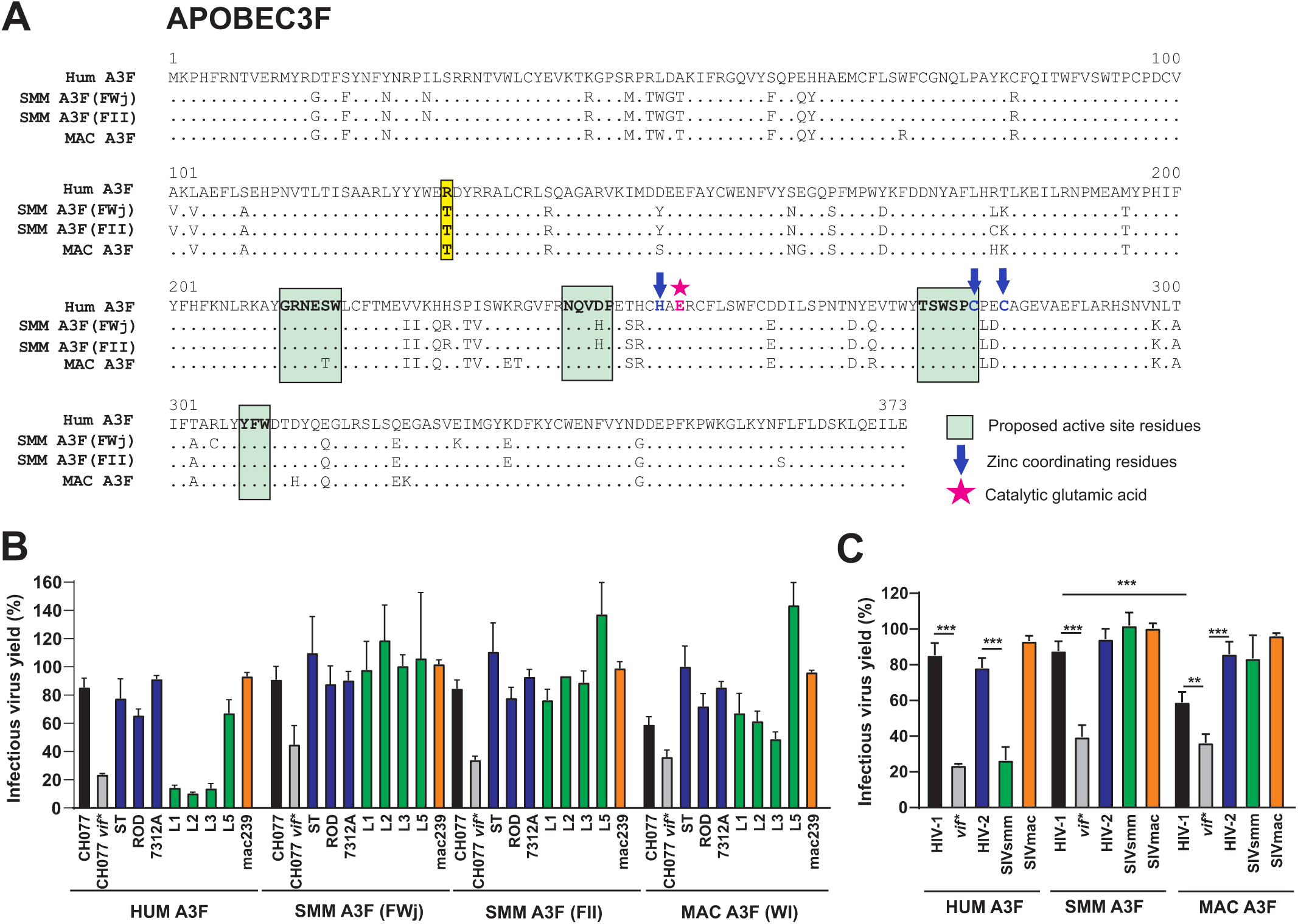
Species-specific effects of human and monkey A3F proteins on HIV-1, HIV-2 and SIVsmm inhibition. (**A**) Amino acid alignment of the human and monkey A3F variants used for functional analysis. Dots indicate amino acid identity and some important domains and residues are highlighted. (**B**) Wild-type and *vif*-defective HIV-1 CH077 and various HIV-2 and SIV proviral constructs and vectors expressing the indicated human, sooty mangabey and rhesus A3F proteins were cotransfected into HEK293T cells and infectious virus yield was measured using the TZM-bl reporter cell infectivity assay. Values were normalized to the infectious virus yield obtained in the absence of APOBEC3 expression construct (100%). Shown is the mean N=4 independent experiments measured in triplicates ± SEM. (**C**) Mean infectious virus yield (±SEM) of the three HIV-2 and four SIVsmm IMCs in the presence of the human or the two SMM or MAC A3F proteins, respectively. HEK293T cells were cotransfected with proviral constructs and APOBEC3 expression vectors and infectious virus yields determined as described in panel A. **, P < 0.01; ***, P < 0.001, calculated using Student’s t test.

For A3G, three macaque orthologues showing variations at 12 amino acid positions and differing in ∼90 residues from the human orthologue were available for analysis (Figure 5A). Most notably, the variation of Y59 to L59/R60 was previously reported to confer resistance to SIVsmm Vif [30]. Predictably, the Vif protein of HIV-1 CH077 efficiently counteracted human A3G (Figure 5B). The three macaque A3G orthologues showed moderate activity against HIV-1 CH077 and were not efficiently counteracted by its Vif protein (Figure 5B, 5C). On average, the SIVsmm strains were slightly more susceptible to the human than to the macaque orthologues, while the opposite was observed for the HIV-2 IMCs (Figure 5C). However, these differences were modest, which agrees with previous data suggesting that A3G does not represent an effective barrier against zoonotic transmission of SIVsmm [29]. SIVsmm L2 was susceptible to the MAC A3G(LR) orthologue but resistant to the remaining two macaque variants (Figure 5B). This agrees with previous data showing that SIVsmm is sensitive to A3G(LR) because its Vif protein contains a Gly at amino acid position 17 and that a G17E substitution renders SIVmac resistant to this A3G variant [30]. The SIVsmm IMCs analyzed in the present study were obtained after passage in macaques (Table S1), which may explain why most of their Vif proteins contained E17 and were resistant to A3G(LR). Notably, HIV-2 strains were less sensitive to inhibition by the macaque A3G(LR) variant than SIVsmm L2 although their Vif proteins generally contain a Gly at position 17.

**Figure 5:**
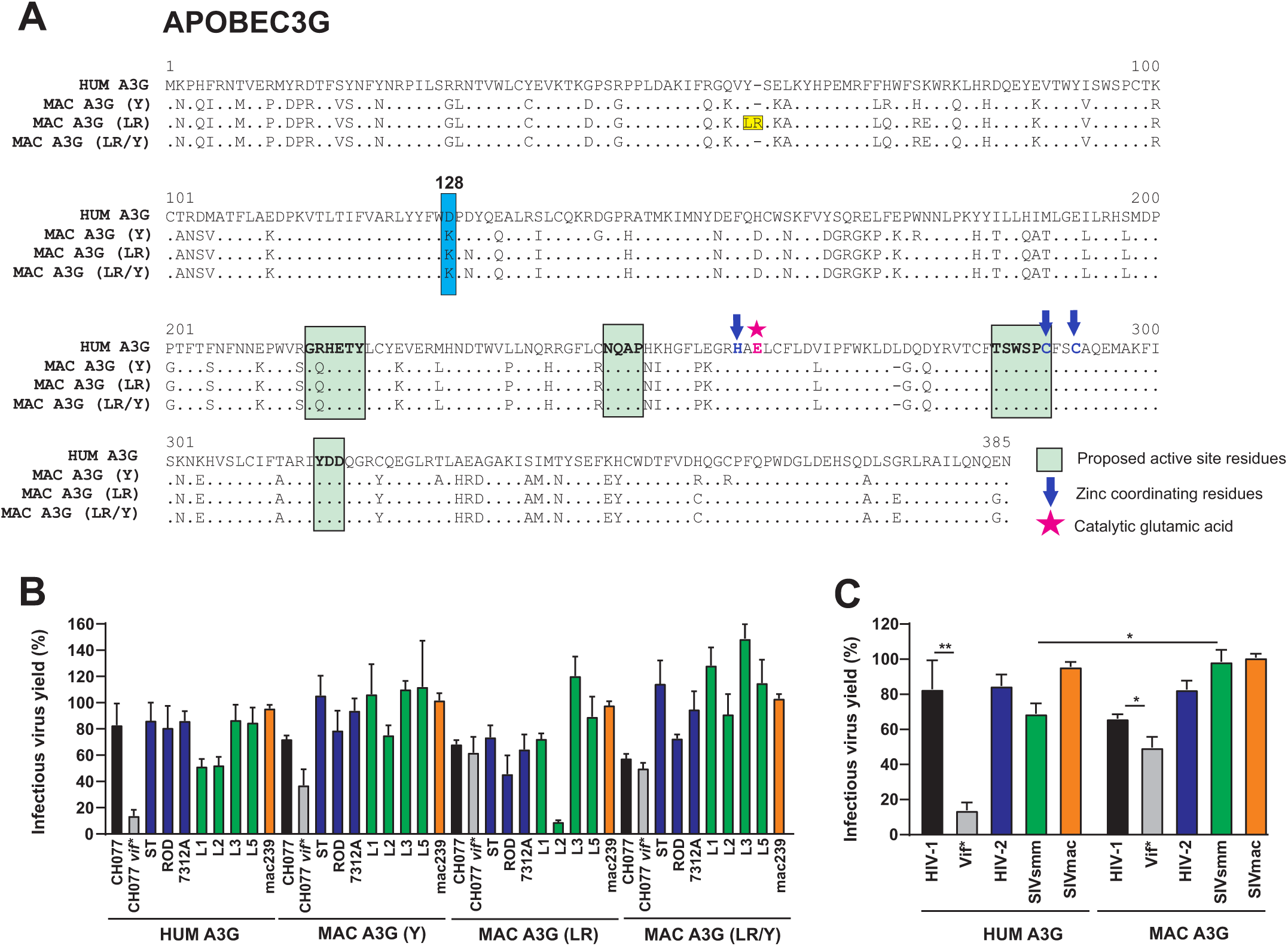
Species-specific effects of human and monkey A3G proteins on HIV-1, HIV-2 and SIVsmm inhibition. (**A**) Amino acid alignment of the human and macaque A3G variants used for functional analysis. Dots indicate amino acid identity and some important domains and residues are highlighted. (**B**) Wild-type and *vif*-defective HIV-1 CH077 and various HIV-2 and SIV proviral constructs and vectors expressing the indicated human and macaque A3G proteins were cotransfected into HEK293T cells and infectious virus yield was measured using the TZM-bl reporter cell infectivity assay. Shown is the mean of N=4 independent experiments measured in triplicates ± SEM. Values were normalized those obtained in the absence of A3F expression construct (100%). (**C**) Mean infectious virus yield (±SEM) of the HIV-2 and SIVsmm IMCs in the presence of the human or MAC A3G proteins, respectively. *, P<0.05; **, P < 0.01, calculated using Student’s t test.

Finally, we examined the susceptibility of the IMCs to inhibition by human and sooty mangabey A3H, which differ in 28 amino acid positions and the presence of the last exon (Figure 6A). In agreement with published data [45], HIV-1 counteracted human but not sooty mangabey A3H (Figure 6B). In contrast, HIV-2, SIVsmm and SIVmac were generally resistant against both human and sooty mangabey A3H (Figure 6B, 6C). Altogether, our results support that A3G and A3H do not represent significant barriers to successful cross-species transmission of SIVsmm to humans. In contrast, human A3F displayed substantially higher activity against SIVsmm than against HIV-2, suggesting specific adaptation to this antiviral factor during viral adaptation to humans.

**Figure 6:**
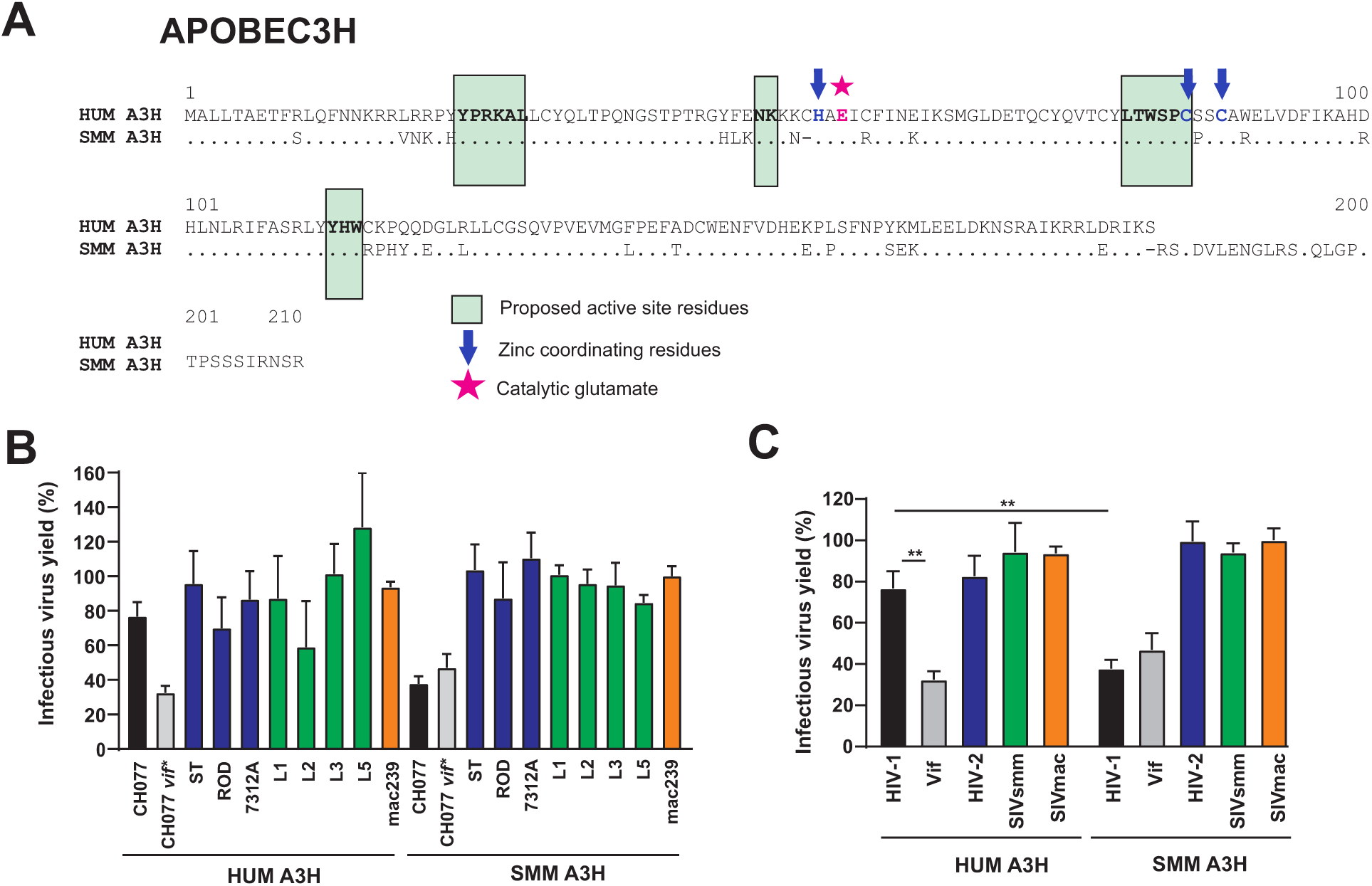
Counteraction of human and sooty mangabey A3H proteins by HIV-1, HIV-2 and SIVsmm. (**A**) Amino acid alignment of the human and SMM A3H variants used for functional analysis. Dots indicate nucleotide identity. Some relevant domains and residues are highlighted. (**B**) Counteraction of the A3H variant shown in panel A by the indicated HIV and SIV IMCs. Shown is the mean N=4 independent experiments measured in triplicates ± SEM. See legend to figure 5 for detail. (**C**) Mean infectious virus yield (±SEM) of the HIV-2 and SIVsmm IMCs in the presence of the human or SMM A3H proteins, respectively. **, P < 0.01, calculated using Student’s t test.

### Role of an R128T variation in A3F-mediated inhibition of HIV-1, HIV-2 and SIVsmm

It has been shown that a single D128K variation in the A3G N-terminal domain plays a key role in the species-specificity of its counteraction by HIV-1 and SIVagm Vif proteins [26,46,47]. Although HIV-1 Vif binds to human A3F in the C-terminal domain on a dispersed interface involving the 11 amino acids from position 255 to 324, it has been reported that HIV-2 Vif interacts with the A3F in the N-terminal domain, specifically amino acids 140-144 [48,49]. However, these residues are generally conserved in A3Fs from human, sooty mangabey, and macaque. In case of A3G and A3H HIV-1, Vif interacts with an amino acid on loop 7, D128 (A3G) or D121 (A3H), respectively. Notably, human A3F differs by a T128R change from sooty mangabey and macaque A3Fs (Figure 4A). To determine whether this loop 7 amino acid substitution contributes to the species-specificity of Vif antagonism, we introduced a R128T mutation in human A3F and the reverse T128R change in sooty mangabey A3F. We found that the R128T change increased the antiviral activity of human A3F against HIV-1 CH077 (Figure 7A) and HIV-2 (Figure 7B), while the reverse change in sooty mangabey A3F had no significant effect on its antiviral activity. Thus, R128T renders human A3F less sensitive to antagonism by HIV-1 and HIV-2 Vif proteins. Human A3F and the T128R mutant sooty mangabey A3F were equally effective against SIVsmm L2 (Figure 7C). In comparison, wildtype sooty mangabey A3F showed the lowest and the R128T human A3F variant the highest activity against SIVsmm L2. Thus, the T128R substitution in sooty mangabey A3F seems to reduce its susceptibility to counteraction by SIVsmm Vif although the differences were modest. Western blot analyses showed that Vif efficiently reduced the levels of A3F in cellular extracts as well as in HIV-1, HIV-2 and (less efficiently) SIVsmm virions (Figure 7, right panels). Our results suggest that *vif*-defective HIV-2 7312 infection results in significant reduction of A3F expression in the cells while this was not observed with *vif*-defective HIV-1 and SIVsmm constructs. This result agrees with our finding that *vif*-defective HIV-2 7312 is less sensitive to A3F and A3G inhibition than HIV-1 and SIVsmm constructs lacking intact *vif* genes (Figure 3). Usually, only modest differences between wildtype and mutant human and sooty mangabey A3F levels were observed in cells transduced with the HIV-1, HIV-2 or SIVsmm proviral construct, although the Western blot confirmed that the R128T substitution increased A3F steady-state expression levels in cells transfected with wildtype HIV-2 7312 (Figure 7B).

**Figure 7:**
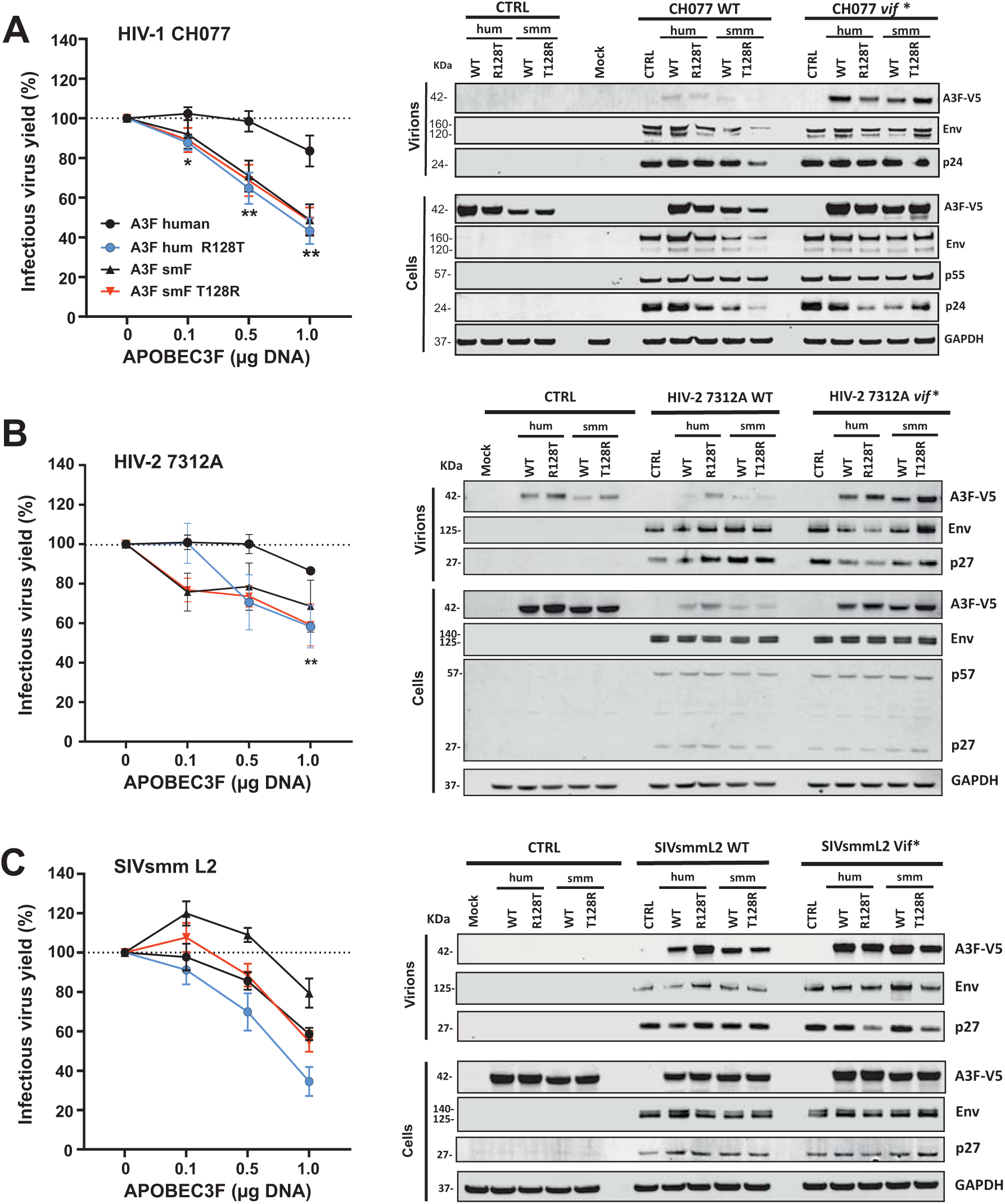
Role of amino acid 128 in restriction of HIV-1, HIV-2 and SIVsmm by A3F. (**A-C**) Proviral constructs of the infectious molecular clones of (A) HIV-1 CH077, (B) HIV-2 7312, or SIVsmm L2 and increasing amounts of plasmid expressing the HA-tagged wildtype and mutant forms of human and SMM A3F were co-transfected into HEK293T cells. Infectious virus yield was measured using the TZM-bl reporter cell infectivity assay. For each proviral construct, values were normalized to the infectious virus yield obtained in the absence of ZAP (100%). Shown is the mean from 3 independent experiments each measured in triplicates ± SEM. The right panels show representative Western blots of HEK293T cells cotransfected with the indicated wildtype or *vif*-defective proviral constructs or an empty control vector (CTRL) and the various A3F expression constructs.

### Changes of Y45H and T84S increase the activity of SIVsmm Vif against human APOBEC3F

To identify variations that might affect the ability of Vif to counteract APOBEC3F in a species-specific manner, we aligned the amino acid sequences of the HIV-2, SIVsmm and SIVmac IMCs analyzed in the present study. In order to focus on those that would be representative of human-specific adaptations, we also compared them and other HIV-2 *vif* sequences found in the database to primary SIVsmm consensus sequence. Nucleotide variation analysis of the HIV-2 *vif* genes, performed using Synonymous Non-synonymous Analysis Program (SNAP), identified multiple positions (shown as red peaks on the upper panel above the alignment in Figure 8A) that were subject to high rates of non-synonymous changes during the evolution of HIV-2. Several of these mutations, such as H28Y, N32R and T84S, showed high, species-specific conservation and were representative of most HIV-2 group A/B and SIVsmm isolates (Figure 8B, S3). These features indicate a likely relevant role in host adaptation of HIV-2 (Figure 8B). Notably, Y28 dominated in both HIV-2 and SIVmac, while most SIVsmm Vif proteins contained H28. Vif proteins of the extremely rare HIV-2 group F, G. H and I strains showed various amino acids at positions 28, 32 and 84 (Figure S3). To determine their functional impact, we introduced substitutions of H28Y and N32R (individually and in combination) into the SIVsmm L2 IMC. In addition, we mutated residue 45 lying in one of the most conserved regions between SIVsmm and HIV-2 (Figure 8A), from Y found in the L2 and L4 Vifs to H present in most SIVsmm and HIV-2 Vif proteins (Figure 8B). We included this change because it is in close proximity to P43H44 reported to play a role in the interaction of Vif with APOBEC3F [50].

**Figure 8:**
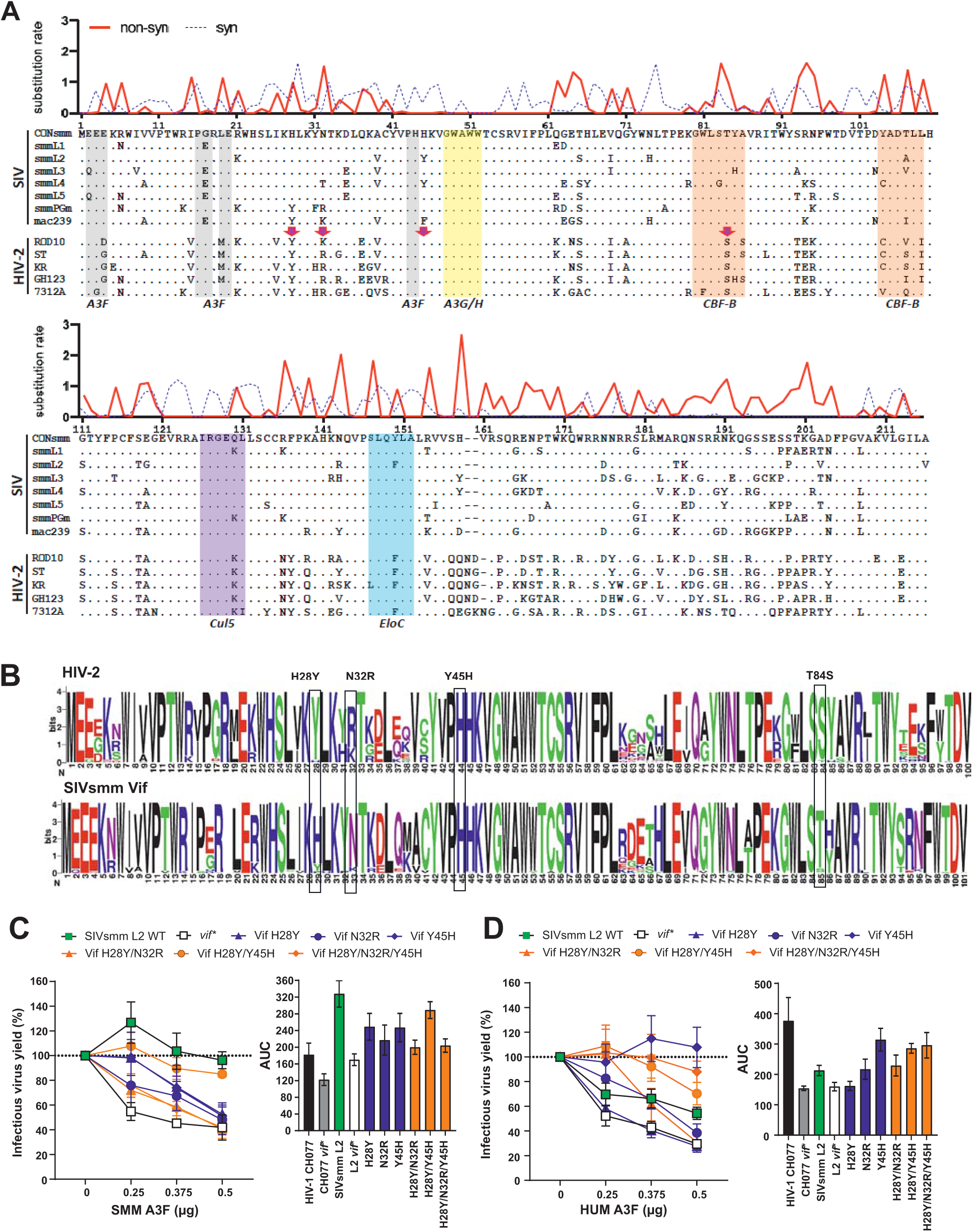
Identification and analysis of potential human-specific adaptation in Vif. (**A**) Alignment of Vif amino acid sequences from HIV-2 and SIV IMCs analyzed in the present study. An SIVsmm consensus Vif sequence is shown on top for comparison. Dots indicate amino acid identity and dashes indicate gaps introduced to optimize the alignment. The upper panel shows HIV-2 synonymous (blue dashed line) and non-synonymous (red line) substitution rates for each codon as compared to SIVsmm consensus Vif. Some known interaction sites and domains are indicated. (**B**) Frequency plots of the N-terminal part of HIV-2 and SIVsmm Vif amino acid sequences. (**C, D**) Effect of mutations predicted to render the SIVsmm Vif more similar to HIV-2 Vif proteins on counteraction of SMM or human A3F. HEK293T cells were cotransfected with the indicated SIVsmm proviral constructs and increasing doses of A3F expression vectors, and infectious virus yield was measured by the TZM-bl reporter assay. Mean infection values (± SEM) were obtained from three or four independent experiments and normalized to those obtained in the absence of A3F expression (100%). The right panels provide AUC (area under the curve) calculated from inhibition curves as shown in the left panels.

For functional analyses, we cotransfected HEK293T cells with the wildtype and mutant proviral SIVsmm L2 constructs and various amounts of sooty mangabey or human A3F expression plasmid and determined infectious virus yield by TZM-bl infection assay (Figure S4). We found that all amino acid changes increased the susceptibility of SIVsmm to sooty mangabey A3F, resulting in a phenotype intermediate between wildtype and *vif*-defective SIVsmm (Figure 8C). In comparison, the Y45H substitution alone or in combination with changes at positions 28 and/or 32 rendered SIVsmm less susceptible to human A3F (Figure 8D). The species-specificity of this effect came as surprise since 45H dominates in both SIVsmm as well as HIV-2 Vif proteins (Figure 8B).

To further determine the effect of the T85S and Y45H changes on the ability of SIVsmm Vif to antagonize A3F, we performed Western blot and infectious virus production assays (Figure 9). We found that both changes increased infectious virus yield in the presence of human A3F from ∼60% to 80% but did not further enhance the already effective counteraction of sooty mangabey A3F (Figure 9A). Western blot analyses indicated that the T85S and Y45H changes do not markedly affect the ability of SIVsmm Vif to induce SMM A3F degradation. For comparison, we used the HIV-2 7312 construct and confirmed that it is hardly susceptible to A3F inhibition, even in the absence of Vif (Figure 9A). Thus, HIV-2 may also have evolved Vif-independent mechanisms to avoid A3F restriction.

**Figure 9:**
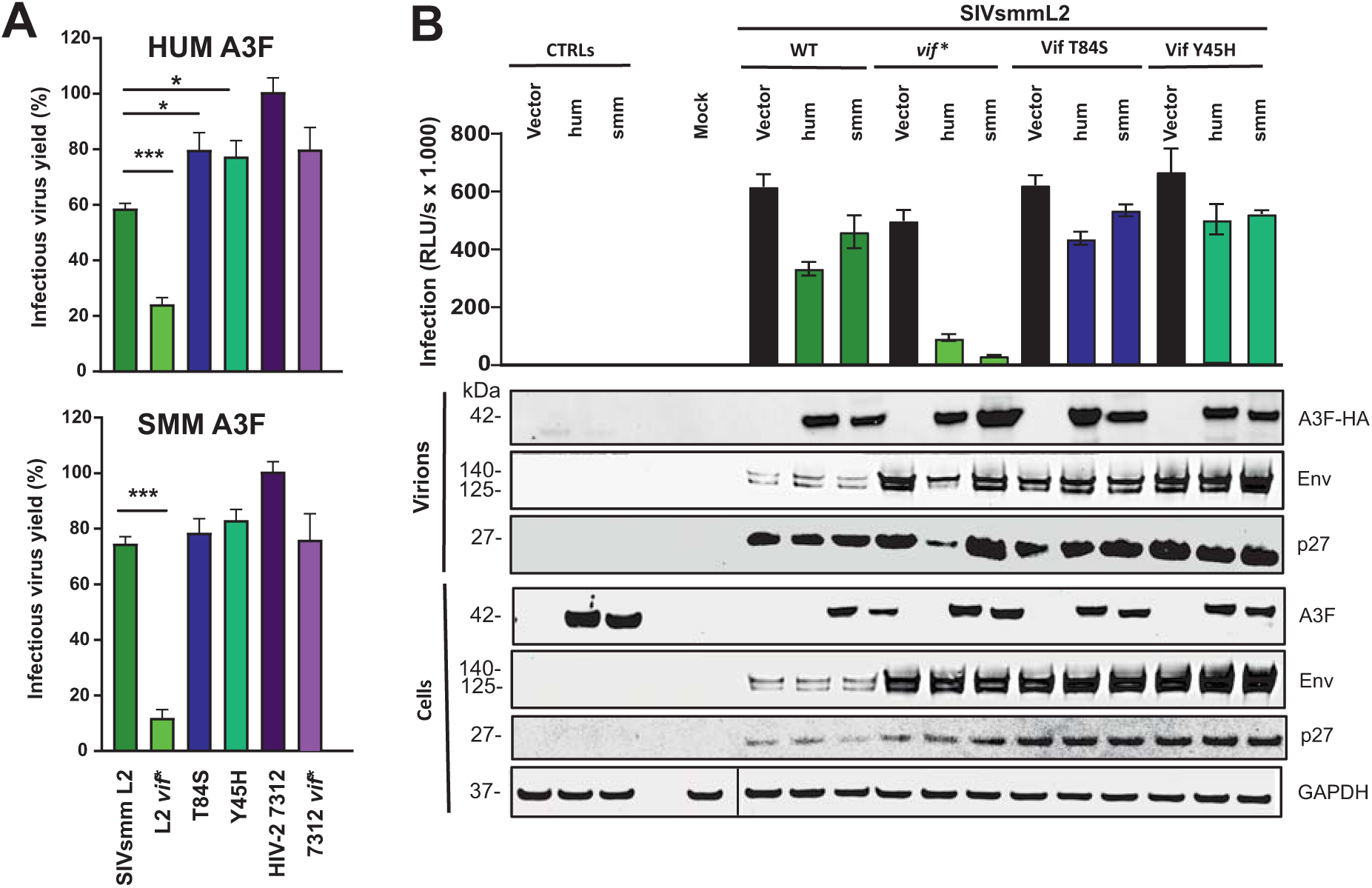
Impact of Y45H and T84S substitutions in Vif on the sensitivity of SIVsmm to inhibition by human or SMM A3F. **(A)** HEK293T cells were cotransfected with the indicated SIVsmm proviral constructs and A3F expression vectors and infectious virus yield was measured by TZM-bl reporter assay. Mean infection values (± SEM) were obtained from three to four independent experiments and normalized to those obtained in the absence of A3F expression (100%). *, P<0.05; ***, P < 0.001, calculated using Student’s t test. (**B**) Infectious HIV-1 yield (top) and expression levels of viral proteins and human or SMM A3F in HEK293T cells transfected with the indicated proviral and A3F expression constructs.

### The T84S change in Vif accelerates SIVsmm replication in human PBMCs

To examine the impact of the identified amino acid variations in Vif on the replication fitness of SIVsmm in human cells, we monitored wt and *vif* mutant infectious virus production in human PBMCs (derived from four different donors) infected with wt and *vif* mutant SIVsmm IMCs over a period of 12 days. For comparison, we included the HIV-2 7312 IMC in the analyses. As expected, the wildtype SIVsmm and HIV-2 IMCs replicated efficiently in human cells whereas the *vif*-defective derivatives did not yield infectious virus (Figure 10A). All five mutant SIVsmm constructs exhibited similar levels of virus production as the wildtype virus. However, the T84S substitution in Vif generally resulted in accelerated replication kinetics (Figure 10B) and slightly higher total virus production (Figure 10C). In comparison, the reverse S84T change did not alter the replicative fitness of HIV-2 7312 in human CD4+ T cells (Figure 10A). Unexpectedly, the H28Y, H28Y/Y45H and H28Y/N32R/Y45H changes that render SIVsmm Vif more similar to HIV-2 Vifs all moderately but significantly reduced infectious virus production in SIVsmm-infected human PBMCs, while the individual Y45H change had no significant effect (Figure 10C). Altogether, these results show that the consistent amino acid differences between SIVsmm and HIV-2 Vif proteins at positions 28 and 32 have a slightly negative effect on viral replication fitness but the T84S change significantly accelerated replication kinetics in primary human PBMCs.

**Figure 10:**
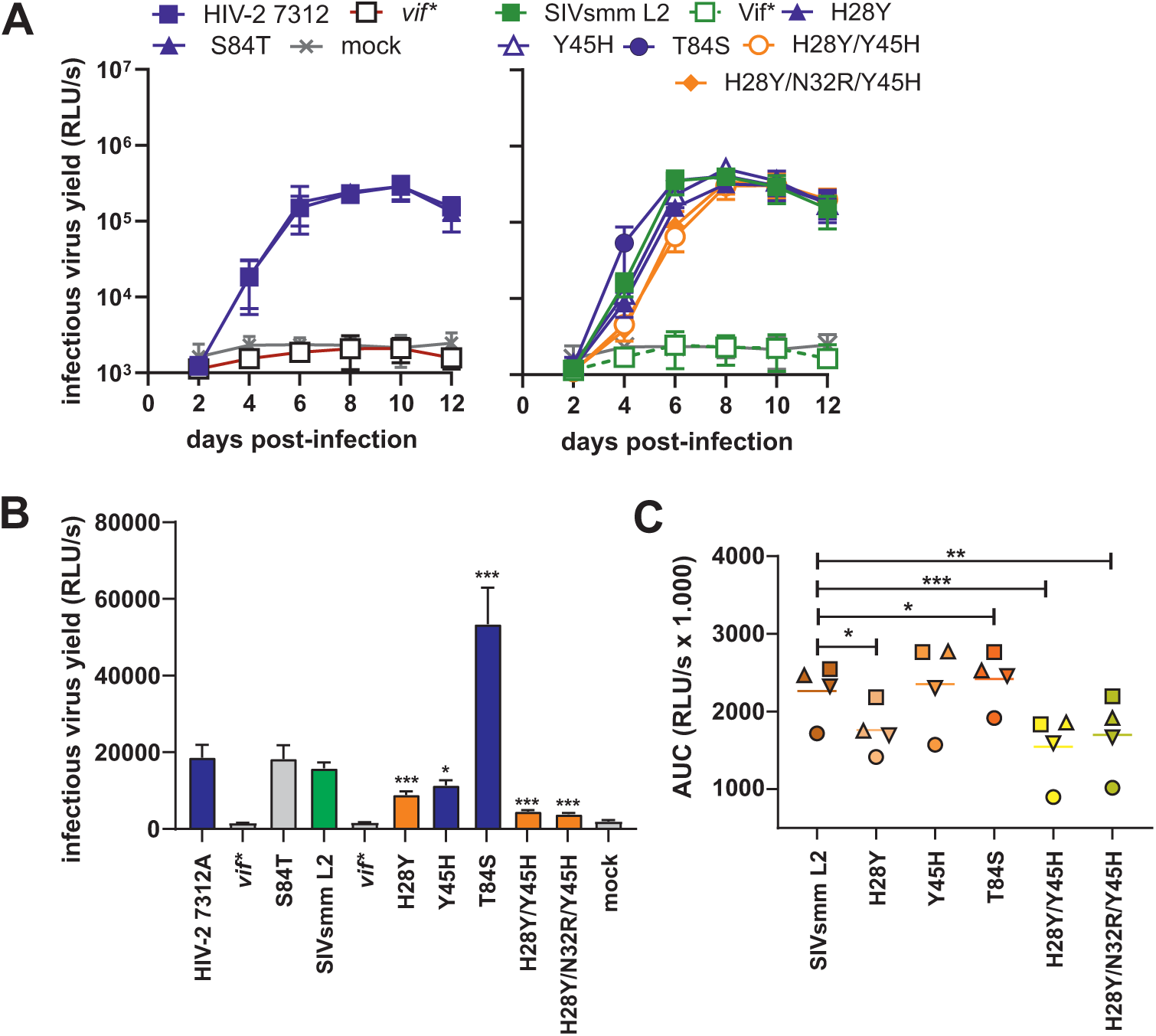
Replication of Vif mutant HIV-2 and SIVsmm IMCs in human CD4+ T cells. **(A)** Mean infectious virus yields measured by triplicate infection of TZM-bl reporter cells with normalized volumes of the supernatants from infected CD4+ T cell cultures derived from four different PBMC donors. (**B**) Infectious virus yields at 4 days post-infection. (**C**) Cumulative infectious virus production in PBMCs infected with the indicated SIVsmm constructs. Shown are AUCs for the replication kinetics as indicated in panel A but calculated for each of the four donors (indicated by a different symbol) individually. *, P<0.05; **, P < 0.01, ***, P<0.001, calculated using Student’s t test for paired comparisons.

## DISCUSSION

It has been shown that primary SIVsmm strains replicate in primary human cells [6] and are capable of counteracting some major human restriction factors, including APOBEC3 proteins without prior adaptation [29]. However, it remained unclear whether HIV-2 acquired increased replication fitness and activity against human APOBEC3 proteins during spread in humans. Here, we show that SIVsmm IMCs differ substantially in their ability to spread in human PBMCs but are, on average, less fit for replication than HIV-2 and (even more) HIV-1. Notably, even those SIVsmm strains showing the highest levels of replication were almost fully inhibited by type I IFN treatment of human PBMCs. Our results clearly show that epidemic HIV-2 group A strains evolved increased ability to counteract or evade the human versions of antiretroviral restriction factors upon zoonotic transmission. Subsequent analyses revealed that the HIV-2 IMCs were on average less sensitive to inhibition by four of the five APOBEC3 family members investigated (i.e. A3D, A3F, A3G and A3H) than SIVsmm. Functional analyses confirmed that HIV-2 Vif proteins degrade human APOBEC3 proteins more efficiently than those of SIVsmm. Thus, our data support that adaptations in Vif that increase its ability to counteract human APOBEC3 proteins were important for the epidemic spread of HIV-2.

Based on previous data using uncloned viral strains [51], we expected HIV-1 to replicate to higher titers in human PBMC than HIV-2 (Figure 1). However, we did not expect that HIV-1 IMCs were on average less efficient in degrading human APOBEC3 proteins than HIV-2 (Figure S2A). Only overexpression of A3H reduced the infectious virus yield of HIV-1 more efficiently than that of HIV-2 (Figure 2B). Altogether, these results agree with previous findings that HIV-1 Vif proteins counteract ABOBEC3 proteins by several degradation-dependent and independent mechanisms [52,53]. It will be interesting to further examine whether the increased ability of HIV-2 to induce degradation of APOBEC3 proteins may compensate for the lack of other counteracting mechanisms exerted by HIV-1.

One limitation of our study is that the HIV-1, HIV-2 and SIVsmm IMCs were generated and obtained in different ways. Except for NL4-3, all HIV-1 IMCs represented TF viruses directly obtained from patient plasma (Table S1). TF HIV-1 strains are usually resistant to the inhibitory effects of IFN-I and efficient in counteracting antiviral restriction factors [33,54]. This explains their relatively efficient replication in the presence of IFN (Figure 1C) and further suggests that the above-mentioned lower activity in degradation of A3 proteins compared to HIV-2 is a common feature of primary HIV-1 strains. With the exception of HIV-2 7312A, all HIV-2 IMCs, as well as two of the seven SIV clones (PG and mac239) were obtained after passage in human cell lines and may not be fully representative of primary HIV-2 and SIVsmm or SIVmac strains. Preadaptation to human cell helps to explain why SIVsmm PG and SIVmac239 replicated efficiently in human PBMCs, at least in the absence of IFN (Figure 1). In contrast, the SIVsmm L1 to L5 IMCs have never been propagated in human cells but were obtained after passage in rhesus macaques (Table 1S) [39,55]. Adaptation to macaques most likely explains why the Vif proteins of most of these SIVsmm IMCs contain a G17E substitution allowing them to counteract the MAC(LR) variant of A3G [30]. The exception was SIVsmm L2, which contains G17 like the Vif proteins of primary sooty mangabey viruses, and was thus selected for mutational analyses. Notably, with the exception of being sensitive to the MAC(LR) variant of A3G, the properties of SIVsmm L2 and its Vif protein were highly similar to those of the remaining four lineages of SIVsmm. Both sooty mangabey and rhesus macaques belong to the *Cercopithecinae* subfamily of Old World monkeys and are genetically closely related. Thus, passage in the latter did most likely not alter most properties of SIVsmm and still allowed meaningful comparison to HIV-2 IMCs.

Four of the five HIV-2 IMC analyzed belong to group A that has spread more efficiently in humans than all remaining groups of HIV-2, albeit much less effectively compared to pandemic HIV-1 group M strains. Notably, some of the consistent amino acid variations between HIV-2 A strains and SIVsmm are shared by group B viruses but not by group F and G HIV-1 Vif sequences available in the Los Alamos HIV sequence data base. Thus, it is tempting to speculate that adaptation to the human host might have been a prerequisite for the epidemic spread of group A and B HIV-2 strains, while the remaining groups, which were usually detected in only single individuals, have adapted less well to their new human host.

Many previous studies have been performed using Vif expression constructs while most experiments performed in the present study involved infectious molecular clones of HIV-1, HIV-2 and SIVsmm. Utilization of highly diverse primate lentiviruses has the limitation that quantitative comparisons are difficult because the viral antigens are highly divergent. However, they are more relevant for the *in vivo* situation and may reveal whether not only Vif function but also other features may contribute to viral susceptibility to inhibition by APOBEC3 proteins. For example, the *vif*-defective HIV-2 7312A IMC was surprisingly resistant to human APOBEC3 proteins (Figure 2B). Our results suggest that different HIV-1, HIV-2 and SIVsmm strains differ substantially in APOBEC3 sensitivity irrespective of Vif function. In addition, our data agree with the evidence that Vif may also counteract APOBEC3 proteins by degradation-independent mechanisms [43,56].

Although four of the seven human APOBEC3 proteins (APOBEC3D, G, F, and H) can restrict replication of HIV-1, APOBEC3G and APOBEC3H are considered to have higher restriction activity than APOBEC3F or APOBEC3D. Surprisingly, for SIVsmm, the APOBEC3F restricted infectious virion production approximately 2-fold more than APOBEC3G, which is opposite to what is observed for HIV-1 [57]. Once an APOBEC3 has bypassed Vif, activity in a virion is determined by APOBEC3 processivity, or ability to deaminate multiple cytosines in a single enzyme/substrate encounter and ability to physically inhibit the reverse transcriptase [58]. In the absence of Vif, APOBEC3G can carry out both activities efficiently, whereas APOBEC3F induces fewer mutations, but can inhibit HIV-1 reverse transcriptase more than APOBEC3G [59,60]. APOBEC3G deamination is also inhibited by physically binding to encapsidated Vif [61,62]. For SIVsmm, the data show that human APOBEC3F was better at resisting Vif-mediated degradation than APOBEC3G so that more was left in cells and thus in virions to exert the restriction activity. Thus, even though biochemically APOBEC3F has been characterized as less active, these activities were largely observed in the absence of Vif. Here, in the presence of Vif, the APOBEC3F was more resistant and thus constituted the main transmission barrier, despite less efficient biochemical activity under ideal conditions.

In the present study, we confirmed that SIVsmm is preadapted for growth in human cells and capable of counteracting human APOBEC3 proteins. We also show, however, the SIVsmm strains that were transmitted to humans were initially most likely highly sensitive to IFN and apparently acquired adaptive changes for epidemic spread in humans, including increased capability to counteract APOBEC3D, F, G and H. We found that a T84S variation in Vif is relevant for species-specific counteraction by HIV-2 and SIVsmm but altogether the determinants seem to be complex. One open question is why HIV-2 A is still less fit for replication and spread in humans (compared to HIV-1) although it crossed the species-barrier about a century ago and its Vif proteins seem to be highly effective at degrading human APOBEC3 proteins.

## Materials and Methods

### Ethical statement

Experiments involving human blood were reviewed and approved by the Institutional Review Board (i.e. the Ethics Committee of Ulm University). Individuals and/or their legal guardians provided written informed consent prior to donating blood. All human-derived samples were anonymized before use. The use of established cell lines (HEK293T, TZM-bl,) did not require the approval of the Institutional Review Board.

### Cell lines

Human Embryonic Kidney (HEK) 293T cells (obtained from the American Type Culture Collection (ATCC) and TZM-bl reporter cells (kindly provided by Drs. Kappes and Wu and Tranzyme Inc. through the NIH AIDS Reagent Program) were cultured in Dulbecco’s Modified Eagle Medium (DMEM) supplemented with 10% heat-inactivated fetal calf serum (FCS), 2 mM L-glutamine, 100 units/ml penicillin and 100 μg/ml streptomycin. TZM-bl cells express CD4, CCR5 and CXCR4 and contain the β-galactosidase genes under the control of the HIV-1 promoter [63,64].

### Expression vectors

Infectious molecular clones of HIV-1, HIV-2 and SIVsmm were described before (summarized in Table S1). Human APOBEC3C, D, F, G and H (haplotype II) expression vectors were obtained from NIH AIDS Reagent and subcloned into a pcDNA expression vector containing a C-terminal HA-tag as reported [57,65]. pcDNA3.2/V5-DEST based macaque and sooty mangabey APOBEC3F constructs containing a C-terminal V5-tag were cloned via the Gateway system. Monkey cDNA was extracted from archival B-cells generated from animals housed at the New England Primate Research Center (NEPRC), and the coding sequences were amplified with primers that introduced an optimal **Kozak sequence** at the 5’ beginning of the <u>APOBEC3F ORF</u> (forward primer: **CAC C**<u>AT GAA GCCTCA CTT CAG AAA CAC AGT GGA GCG AAT G</u>; reverse primer: CTC GAG AAT CTC CTG CAG CTT GC). Rhesus macaque A3G and sooty mangabey A3H expression constructs have been described [30,66].

Generation of mutant HIV and SIV IMCs. The Vif-deficient versions of HIV-1 NL4-3, HIV-1 CH077, HIV-1 CH058, HIV-2 AB 7312A, SIVsmm L2. RM136, SIVsmm L5.DE28 and SIVmac239 IMCs were generated by introducing 2 stop codons directly after the end of the *pol/vif* overlapping reading frame. Mutations inducing amino acid substitutions in the *vif* region of infectious molecular clones or in the APOBEC3 expression constructs were introduced using Q5 site-directed mutagenesis kit (NEB). The presence of the desired mutations was confirmed by Sanger sequencing.

### PBMC isolation and stimulation

Peripheral blood mononuclear cells (PBMCs) from healthy human donors were isolated using lymphocyte separation medium (Biocoll separating solution; Biochrom). Cells were cultured in RPMI 1640 medium supplemented with 10% FCS, 2 mM glutamine, streptomycin/penicillin (Gibco) and IL-2 (10 ng/ml) (Miltenyi Biotec). PBMCs were stimulated for 3 days with 2 μg/ml PHA and if indicated, with 500u/ml IFNα (PBL Assay Science) starting 24h before infection.

### Virus stock preparation

To generate virus stocks, HEK293T cells were transfected with the proviral HIV or SIV DNA (5µg per well of a 6-well plate) using calcium phosphate method. Two days post-transfection, supernatants containing infectious virus were harvested. To normalize the amount of virus dose, elative infectivity was measured by TZM-bl assay and the activity of viral reverse transcriptase present in the supernatant was measured as described before [67].

### Viral replication kinetics

Pre-stimulated PBMCs of different donors were infected with normalized virus stocks. The following day, cells were washed 3x in PBS to remove input virus and each sample was split into triplicates and transferred to a 96-well plate. Virus-containing supernatant was harvested every 2 days and fresh medium RPMI 1640 medium supplemented with 10% FCS, 2 mM glutamine, streptomycin/penicillin (Gibco) and IL-2 (10 ng/ml) (with or without 500u/ml IFNα) was added, up until day 10 or 12 (as indicated). The relative amount of infectious virus present in the harvested supernatants was measured by TZM-bl cell-based infectivity assay.

### Infectivity assay

TZM-bl cells (10.000/well) were seeded in 96-well plates and infected with equal volumes of virus-containing cell supernatants. Infections were performed in duplicate (viral kinetics) or triplicate (transfections). Two days post-infection, viral infectivity was detected using the Gal-Screen kit from Applied Biosystems as recommended by the manufacturer. β-galactosidase activity was quantified as relative light units per second using microplate luminometer (Orion).

### Lentiviral susceptibility to human APOBEC3 proteins

HEK293T cells were co-transfected with 4:1 ratio of proviral DNA and pcDNA APOBEC3 expression construct or pcDNA empty vector. Virus containing supernatants were harvested 2 days later and the produced infectious virus yield was measured using TZM-bl reporter cell line and Gal-Screen kit from Applied Biosystems as recommended by the manufacturer. β-galactosidase activity was quantified as relative light units per second using the Orion microplate luminometer.

### Lentiviral susceptibility to simian APOBEC proteins

To assess relative lentiviral sensitivity to APOBEC proteins overexpression, HEK293T cells (in 24-well format) were co-transfected using PEI transfection reagent with 0.75 µg of the indicated IMCs and 0.25 µg of pcDNA3/V5-DEST APOBEC3 expression constructs. Virus containing supernatants were harvested 2 days later and used to infect TZM-bl reporter cells in triplicates. β-galactosidase activity was measured 2 days later using Gal-Screen kit (Applied Biosystems) as relative light units per second using microplate luminometer. Infectious virus yield values of each IMC in the presence of A3 proteins were normalised to their corresponding pcDNA3/V5-DEST empty vector only control.

### Western blot

Cells were co-transfected in 12-well plates with 2 µg of indicated IMC and 0.5 µg DNA of pcDNA3/V5-DEST APOBEC3 expression vector or empty pcDNA3/V5-DEST vector. Two days post-transfection, cells were lysed with Co-IP buffer (150 mM NaCl, 50 mM HEPES, 5mM EDTA, 0.10% NP40, 0.5 mM sodium orthovanadate, 0.5 mM NaF, protease inhibitor cocktail from Roche) and cell-free virions were purified by centrifugation of cell culture supernatants through a 20% sucrose cushion at 20,800 g for 90 minutes at 4°C and lysed in CO-IP lysis buffer. Samples were reduced in the presence of β-mercaptoethanol by boiling at 95°C for 10 min. Proteins were separated in 4 to 12% Bis-Tris gradient acrylamide gels (Invitrogen), blotted onto polyvinylidene difluoride (PVDF) membrane, and incubated with anti-V5(Cat#13202; Cell Signaling, 1:5000), anti-HIV-1 Env (Cat #12559, obtained through the NIH AIDS Reagent Program, 1:1000) or anti-HIV-2 Env (Cat #771, obtained through the NIH AIDS Reagent Program, 1:500), anti-p24 (Cat #ab9071; Abcam, 1:1000) and or anti-p27(Cat#2321, obtained through the NIH AIDS Reagent Program 1:200), anti-GAPDH (Cat #607902, Biolegend, 1:1000). Blots were probed with IRDye® 680RD Goat anti-Rabbit IgG (H + L) (Cat #926-68071, LI-COR), IRDye® 800RD Goat anti-Mouse IgG (H + L) (Cat #926-32210; LI-COR) and IRDye® 800RD Goat anti-Rat IgG (H + L) (CAT#925-32219; LI-COR) Odyssey antibodies all diluted to 1:20000 and scanned using a LI-COR Odyssey reader.

### A3 degradation assay

To compare efficiency of different Vif in mediating degradation of A3G-HA and A3F-HA, 293T cells (1 × 10^5^ per well) in 12 well plates were co-transfected with either A3G-HA (500 ng) or A3F-HA (500 ng or 100 ng) and titration of pCG-Vif-AU1 expression plasmids (0, 25, 50, 100 and 200 ng) using GeneJuice (Novagen) transfection reagent. To equalize the amount of plasmid DNA transfected, empty pCG-AU1 vector was used. Then, 24 h post transfection the media was changed. After 44 h post transfection, cells were washed with PBS and lysed using 2× Laemmli buffer. Total protein in the cell lysate was estimated using the Lowry assay and 30 µg total protein from each cell lysate was used for immunoblotting with anti-HA antibody (Cat# H9658, Sigma, 1:10000), anti-AU1 antibody (Cat# ab3401, Abcam, 1:1000) and anti-α tubulin antibody (Cat # PA1-20988, Invitrogen, 1:1000) as primary antibodies. Secondary detection was performed using Licor IRDye antibodies produced in goat (IRDye 680-labeled anti-rabbit 1:10000 Cat # 926-68071 and IRDye 800-labeled anti mouse 1:10000 Cat # 926-32210). Immunoblots were quantified using Image Studio Software with normalization of each experimental lane to its respective α-tubulin band.

### HIV-2 and SIVsmm sequence analysis

SIVsmm and HIV-2 isolate *vif* sequences were obtained from Los Alamos HIV sequence database (www.hiv.lanl.gov). Consensus SIVsmm sequence was generated using available Vif amino acid *s*equences of SIVsmm primary isolates and Consensus Maker online tool (https://www.hiv.lanl.gov/content/sequence/CONSENSUS/consensus.html). To compare substitution ratios among available HIV-2 *vifs* in reference to SIVsmm consensus, nucleotide sequences were aligned using Clustal Omega [68] and analysed using SNAP v2.1.1 (https://www.hiv.lanl.gov/content/sequence/SNAP/SNAP.html?sample_input=1). Average synonymous and non-synonymous substitution changes in each codon were plotted in Prism.

### Statistical analysis

Statistical analysis was performed using GraphPad Prism software. Two-tailed Student’s *t*-test was used to determine statistical significance. Unpaired tests were used with the exception of experiments involving PBMCs obtained from different donors. Significant differences are indicated as follows: *, *P* < 0.05; **, *P* < 0.01; ***, *P* < 0.001; ****, *P* < 0.0001.

## Acknowledgments

We thank Martha Mayer, Kerstin Regensburger and Regina Burger for technical assistance and Michael Emerman and Molly Ohainle for providing A3H constructs. TZM-bl cells were obtained through the NIH AIDS Reagent Program, Division of AIDS, NIAID, NIH: TZM-bl from Dr. John C. Kappes, Dr. Xiaoyun Wu and Tranzyme Inc. This work was funded in part by NIH AI083118 (W.E.J.) and R01 AI050529 and UM1 AI 126620 (B.H.H.).

## LEGENDS TO SUPPLEMENTAL FIGURES

**Figure S1.**
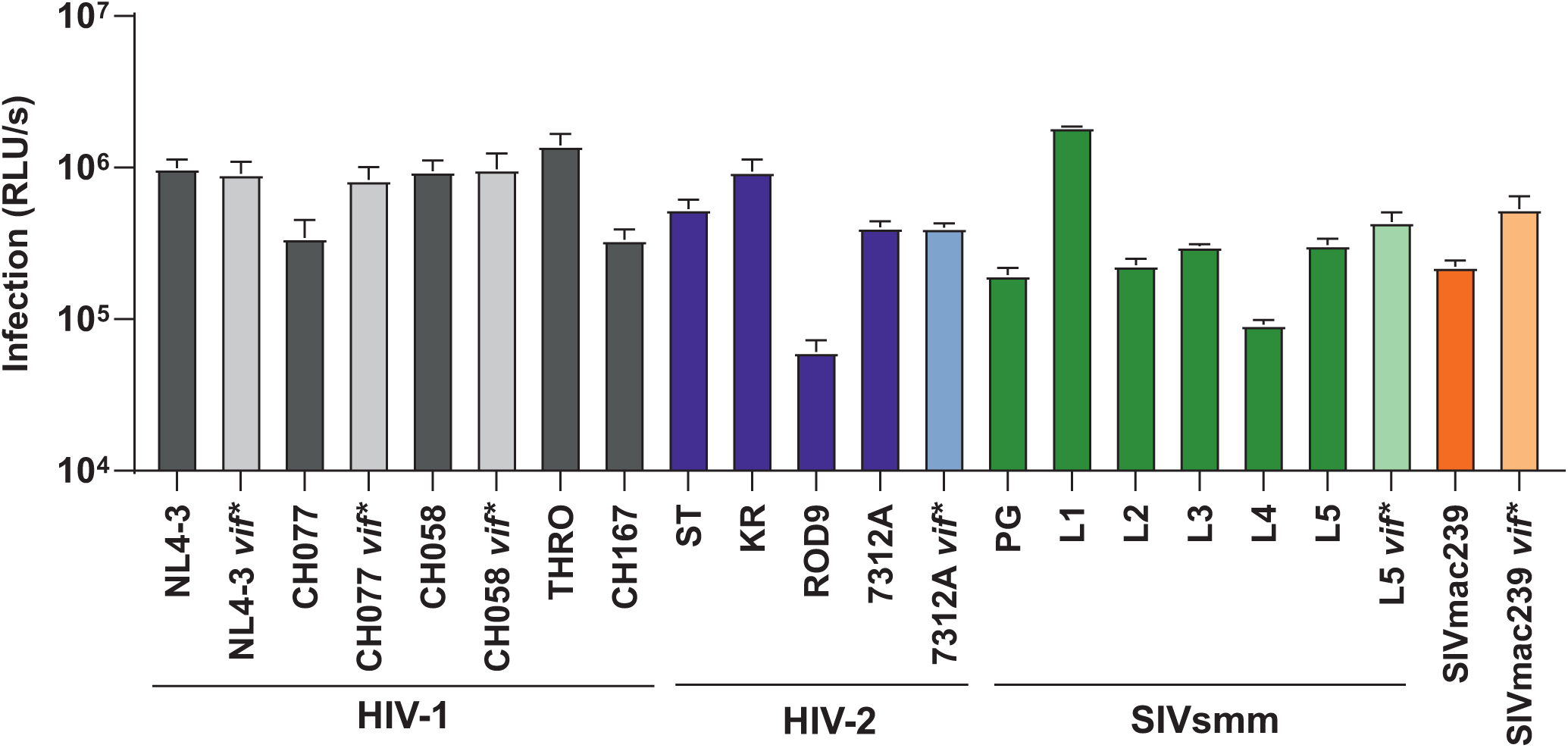
Virus production by wild-type and *vif*-defective HIV-1, HIV-2 and SIV constructs in the absence and presence of transient expression of the indicated APOBEC3 proteins. HEK293T were cotransfected with the indicated HIV and SIV proviral constructs and expression vectors for the indicated A3 proteins or empty vector (CTRL). Supernatants were harvested 2 days later, and infectious virus yield was determined by infecting TZM-bl indicator cells. Shown are average values of a representative experiment measured in triplicates ±SD.

**Figure S2.**
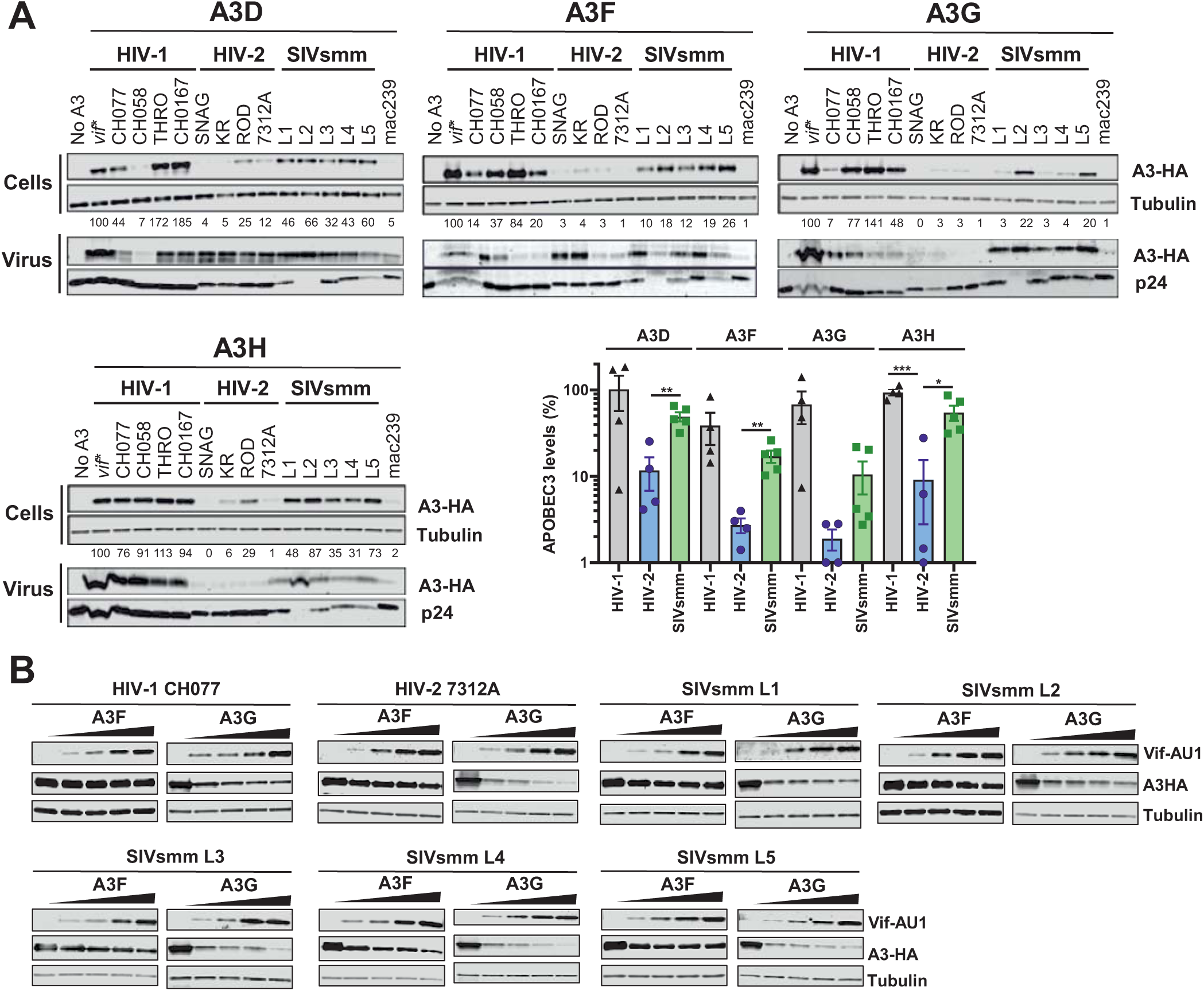
Sensitivity of HIV-1, HIV-2 and SIVsmm to human APOBEC3-mediated restriction. (A) Proviral constructs of indicated IMCs of HIV-1, HIV-2 or SIVsmm and a plasmid expressing the indicated HA-tagged APOBEC3 protein were cotransfected into HEK293T cells. Viral supernatant and cell lysates were collected 48 hours post-transfection. Proteins from cell lysate and viral lysate were separated by SDS PAGE, transferred to nitrocellulose membrane and probed using antibodies, as labeled, with α-tubulin or HIV-1 p24 serving as the loading controls. The HIV-1 p24 antibody was not able to cross react and detect the p27 capsid protein from all SIVsmm IMC. Thus, only cell lysate immunoblots were quantified with normalization of each experimental lane to its respective α-tubulin (bar graph). (B) Vif mediated degradation of A3F and A3G. 293T cells were cotransfected with constant quantity of A3F or A3G expression plasmid (500ng) and increasing amounts of HIV or SIV Vif –AU1 expression constructs. Cell lysate was collected 48 hours and proteins from cell lysates were separated by SDS PAGE, transferred to nitrocellulose membrane and probed using specific antibodies, as labeled, with α-tubulin serving as the loading control.

**Figure S3.**
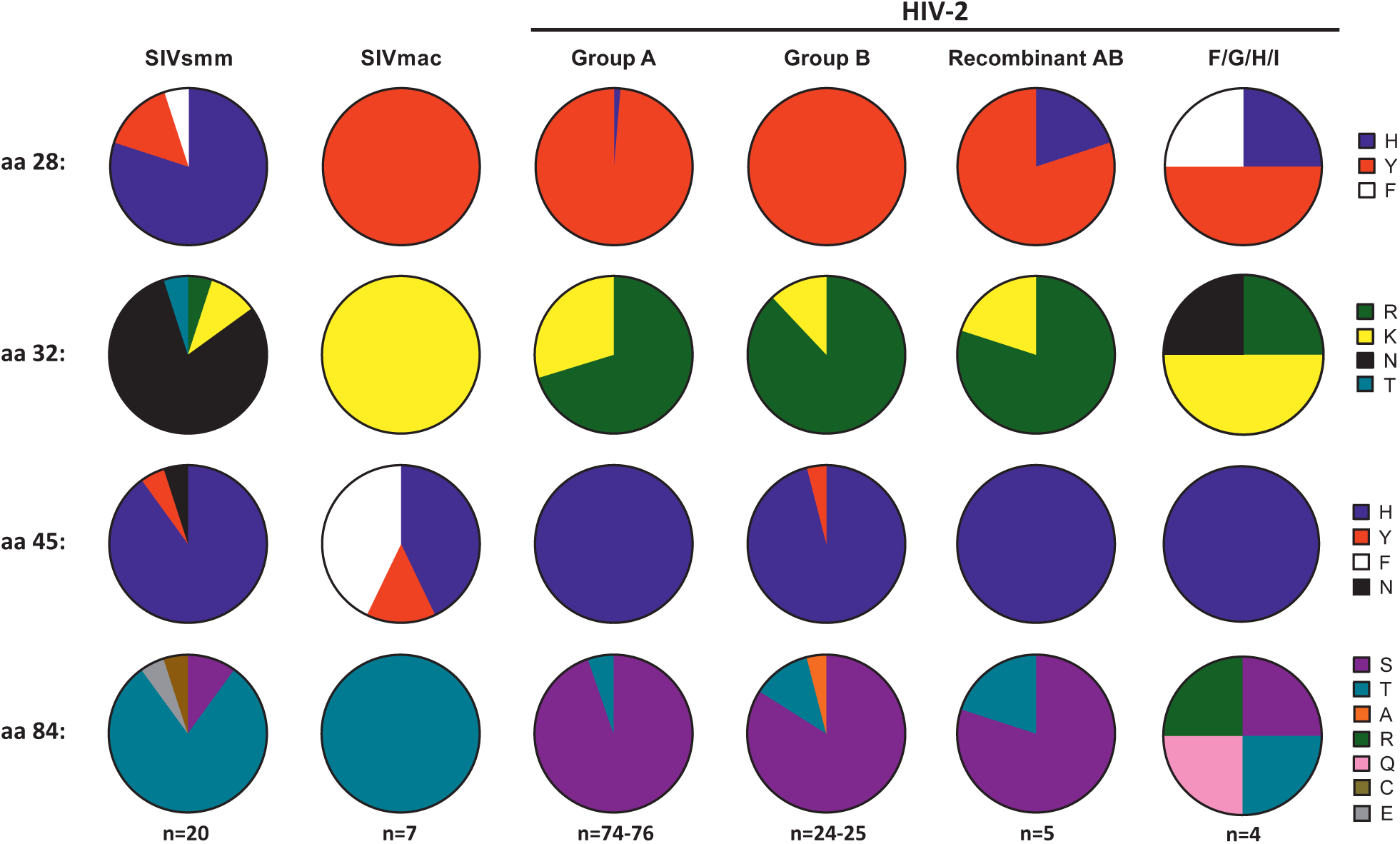
Frequency of specific amino acid residues at positions 28, 32, 45 and 84 in SIV and HIV-2 Vif proteins. Amino acid (aa) frequencies were calculated based on the 2018 Los Alamos HIV-2/SIVsmm Vif protein curated alignment. Number of sequences may differ because in some cases the amino acid was not known.

**Figure S4.**
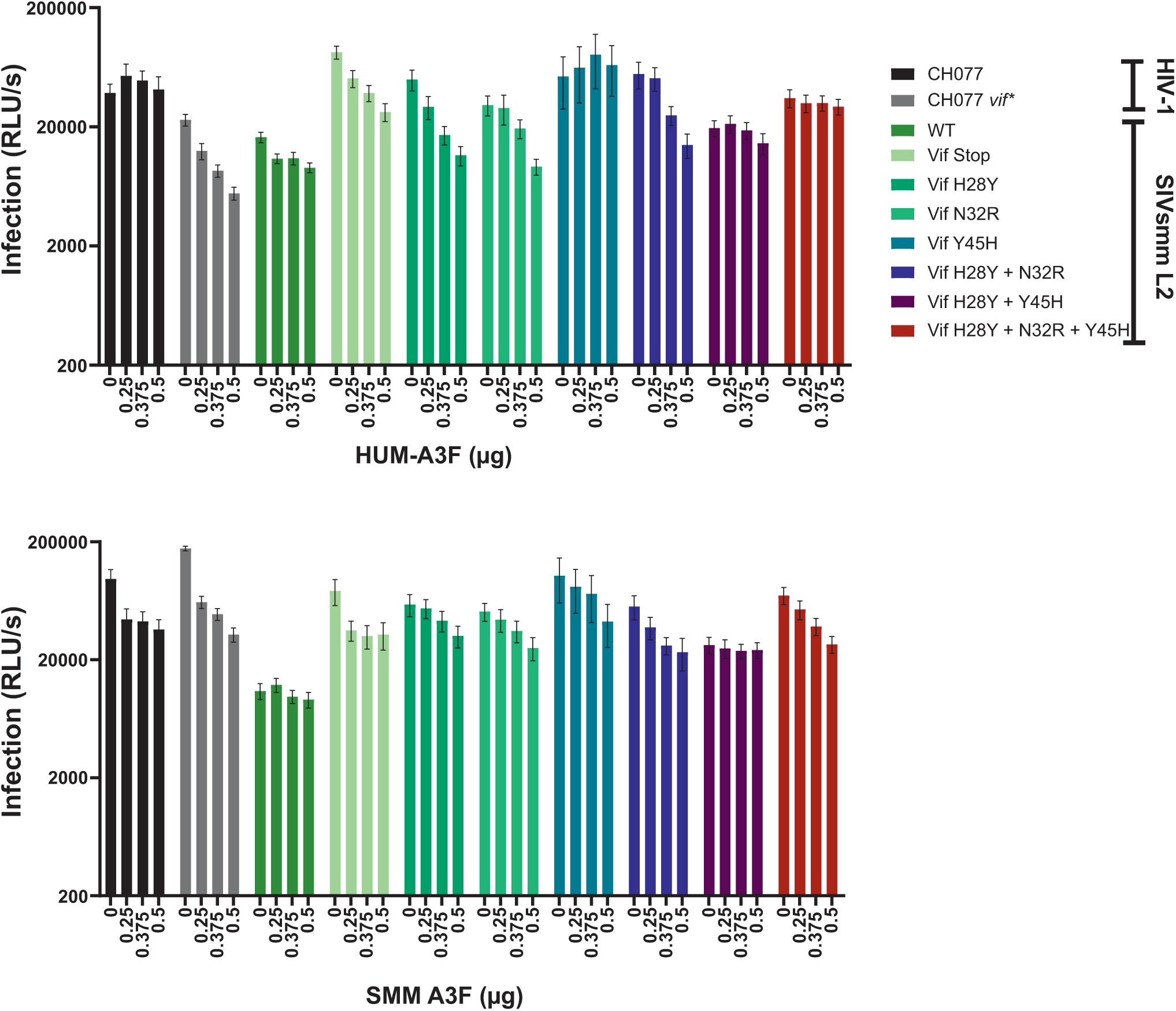
Virus production by wild-type and Vif mutant HIV-1and SIVsmm constructs in the absence and presence of increasing amounts of human (upper) or SMM (low) A3F proteins. Proviral constructs of the indicated infectious molecular clones of HIV-1 or SIVsmm L2 and increasing amounts of a plasmids expressing the hum or smFII A3F were co-transfected into HEK293T cells, the amount of DNA was kept constant by addition of an empty vector. Infectious virus yield was measured using TZM-bl reporter cell infectivity assay. Shown are the mean values from at least 3 independent experiments each measured in triplicates ± SEM.

**Table S1.** Features and origin of HIV and SIV strains examined.

| Virus | Group/Subtype or lineage | IMC | Name used | Origin | References |
| --- | --- | --- | --- | --- | --- |
| HIV-1 | M/B | NL4-3 | HIV-1 M B NL4-3 | Laboratory generated recombinant of HIV-1 isolates NY5 and BRU from symptomatic, chronically infected MSMs; virus was passaged in T cell lines prior to isolation | [1] |
|  | M/B | CH77_TF1 | HIV-1 M B CH077 | MSM patient diagnosed with acute HIV-1 infection; directly isolated from patient's plasma | [2] |
|  | M/B | CH58_TF1 | HIV-1 M B CH058 | MSM patient diagnosed with acute HIV-1 infection, directly isolated from patient's plasma | [2] |
|  | M/B | THRO_TF1 | HIV-1 M B THRO | MSM patient diagnosed with acute HIV-1 infection, directly isolated from patient's plasma | [2] |
|  | M/C | CH167 | HIV-1 M C CH167 | 28-year old, sexually infected female patient diagnosed with chronic HIV-1 infection, directly isolated from patient's plasma | [3] |
| HIV-2 | A | ROD | HIV-2 A ROD9/10 | 32-year old male AIDS patient; virus was isolated following propagation in T cell line | [4] |
|  |  | HIV2ST JSP4-27 | HIV-2 A ST | Asymptomatic female sex worker, virus isolated following propagation of patient's PBMCs with HUT-78 cells | [5,6] |
|  |  | GH123 | HIV-2 A GH123 | Symptomatic 33-year old female sex worker, virus isolated following propagation of patient's PBMCs with MOLT-4/8 T cells; GH123 is derived from isolate GH1 | [7–9] |
|  |  | KR | HIV-2 A KR | anonymous patient, virus isolated following following propagation of patient's PBMCs with MOLT-4/8 T cells | [10,11] |
|  | A-B recombinant | 7312A | HIV-2 AB 7312A | Symptomatic 32-year old male patient, virus isolated following co-culture of PBMCs with uninfected PBMCs | [12] |
| SIVsmm | lineage 1 | SIV.L1. RM174.tf.v1 | SIVsmmL1 | Rhesus macaque experimentally infected with SIVsmm; virus was passaged in two macaques before isolation | [13,14] |
|  | lineage 2 | SIV.L2 RM136.tf | SIVsmmL2 | Rhesus macaque exp. infected with SIVsmm isolate from naturally infected 16-year old male sooty mangabey; virus was passaged three macaques before isolation | [13,14] |
|  | lineage 3 | SIV.L3 RM175.tf | SIVsmmL3 | Rhesus macaque exp. infected with an SIVsmm isolate from a naturally infected 14-year old male sooty mangabey; virus was passaged in two macaques before isolation | [13,14] |
|  | lineage 4 | SIV.L4 RM57.tf | SIVsmmL4 | Rhesus macaque exp. infected with SIVsmm isolate from naturally infected 30-year old female sooty mangabey; virus was passaged in two macaques before isolation | [13,14] |
|  | lineage 5 | SIV.L5 DE28.tf | SIVsmmL5 | Rhesus macaque exp. infected with SIVsmm from 13-year old male sooty mangabey; virus was passaged in two macaques before isolation | [13,14] |
|  | n.a. | PGm5.3 | SIVsmm PG | Pig-tailed macaque exp. infected with SM blood containing SIVsmm; virus was passaged in human PBMCs before isolation | [15] |
| SIVmac | n.a. | 239 | SIVmac239 | Rhesus macaque exp. infected with extensively rhesus-passaged virus of SM origin; isolated following passage in HUT-78 cells | [16,17] |

